# Enhancing breakpoint resolution with deep segmentation model: a general refinement method for read-depth based structural variant callers

**DOI:** 10.1101/503649

**Authors:** Yao-zhong Zhang, Seiya Imoto, Satoru Miyano, Rui Yamaguchi

## Abstract

**Motivation:** For short-read sequencing, read-depth based structural variant (SV) callers are difficult to find single-nucleotide-resolution breakpoints due to the bin-size limitation.

**Results:** In this paper, we present RDBKE to enhance the breakpoint resolution of read-depth SV callers using deep segmentation model UNet. We show that UNet can be trained with a small amount of data and applied for breakpoint enhancement both in-sample and cross-sample. On both simulation and real data, RDBKE significantly increases the number of SVs with more precise breakpoints.

**Availability:** source code of RDBKE is available at https://github.com/yaozhong/deepIntraSV

## 1 Background

Structural variants (SVs) are genomic alterations with a length that at least 50 base pairs (bp). When compared with smaller size mutations, such as single nucleotide polymorphisms (SNPs) and indels, the larger size of SVs makes them more likely to alter genomic structures and have functional consequences. In many diseases, such as Alzheimer’s disease and cancer, SVs have been found to play important roles [1, 2, 3, 4]. With the recent progress of sequencing technologies, more accurate and comprehensively detection of SVs in the whole genome-scale becomes possible.

To detect SVs on NGS data, many different algorithms have been developed. Those methods utilize different types of information derived from NGS reads. Generally, those methods can be categorized into four major types: *read depth (RD)* [5], *paired-end mapping (PEM)* [6], *split-read (SR)* [7] and *assembly-based (AS)* [8]. An **RD** method first divides the whole genome into non-overlapping bins and calculates averaged RD for each bin. Duplication (DUP) and deletion (DEL) are detected by finding abnormal RD changes for adjacent bins. **PEM** makes use of the span and orientation of paired-end reads, which can detect SVs, such as inversion. Note that it is difficult for RD and PEM to find nucleotide-resolution breakpoints. If an adjoining short-read is separately aligned in different coordinates of a reference genome, we can make use of **SR** information to determine SVs and breakpoints in nucleotide-resolution. A more universal approach is AS that assembly short-read into a draft genome and compared it with the reference genome for search potential SVs. But assembly-read is usually computational cost for short-read sequencing. Besides the above four types, several methods integrate more than one type of information or results from different methods. For example, DELLY [6] integrated both PE and SR for SV detection. Recently, with accurate profiling of SVs becomes possible, several works made systematic comparisons for the current SV detection algorithm and make benchmarks for evaluating new SV detection algorithm. Kosugi et al. [9] compared 69 SV detection algorithms for short-reading whole-genome sequencing (WGS) data. For the RD-based, bin-size is an important parameter that affects SV detection performance. On one hand, a larger bin-size is more reliable to capture larger SVs, such as CNVs, at the cost of sensitivity loss for small-length SVs. On the other hand, data based on small bin-size is usually very noisy, and surrounding SNPs or indels make the SV prediction even difficult.

In this paper, we focus on the RD-based SV detection approach and develop a deep segmentation model to enhance the breakpoint resolution of SVs through further analysis surrounding single-nucleotide RD information. The method is motivated to exceed the bin size limitation of a general RD-base SV-caller and make it possible to predict single-nucleotide-resolution breakpoints for RD data. Compared with recent proposed deep learning methods for SV related tasks, our method has the following novel points: **first, formalizing the breakpoint detection problem as a segment task.** Usually, deep learning models are applied as a classifier. For example, deepVariant [10] uses a well-calibrated convolutional network to call SNP and indels (≤ 50 bp) categories (homozygous reference, heterozygous and homozygous alternate) based on image pileups around putative sites. Instead of only classifying whether breakpoints exist inside of a region, we segment SV components inside of the putative region and infer breakpoints in single-nucleotide-resolution. **Second, the proposed method is integrated seamlessly with existing RD methods, which leverages the power of the traditional approach and deep learning models.** We use a traditional RD method to initially detect bin-resolution breakpoints, and use the proposed segmentation model to scan nucleotide-resolution RD information surrounding the breakpoints and make the further refinement. We conduct a series of experiments on both simulation and real WGS data. For the real data, we perform systematic experiments for in-sample and cross-sample applications.

## 2 Methods

### 2.1 Overall breakpoint enhancement pipeline

Figure 1(a) introduces the pipeline of RDBKE for enhancing the breakpoint of an RD-based SV-caller in single-nucleotide resolution. Before enhancement, an RD-based SV-caller is first applied to get initial SV predictions. Without loss of generality, we take the commonly used CNVnator [5] for example. In a typical RD-based SV caller, reads are first aligned to a reference genome and the reference genome is split into non-overlapping bins with a fixed length. For each bin, the number of mapped reads covered by the bin is counted as the smoothed RD signal. The bin-size for an RD-based SV caller is a very important parameter, which determines the sensitivity of predictions and breakpoint resolution. Considering a further step enhancement, a smaller bin-size (e.g., 50 bp) is preferred to increase the number of candidate SVs predicted by the original SV caller. In the enhancement stage, we zoom into the genomic regions surrounding breakpoints in a fixed-length window and apply a trained deep segmentation model to mask SV associated coordinates according to the base-wise RD information in the screening window. The new breakpoints are inferred based on the SV masks and updated accordingly, shown in Figure 1(b).

**Fig. 1:**
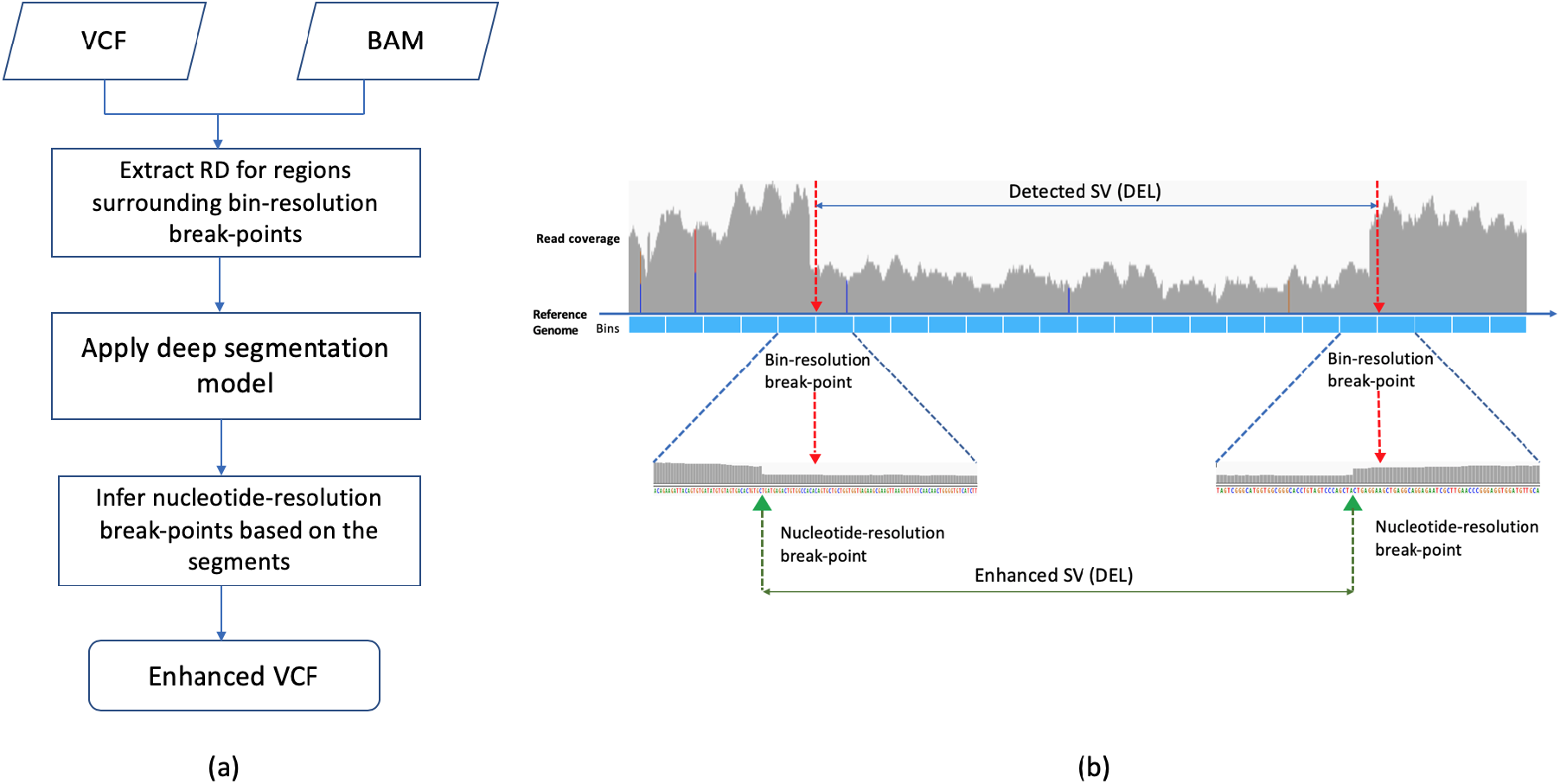
The overall pipeline of RDBKE, that enhances an RD-based SV caller with deep segmentation model. (a). The overall workflow of the enhancement for a general RD-based SV-caller. (b). An example illustrates enhancing the original bin-resolution breakpoints using the RD information surrounding the breakpoint candidates.

### 2.2 UNet segmentation model

Generally speaking, the base-wise read-depth information is too noisy to be directly processed. The key motivation of using the deep segmentation model for breakpoint enhancement is **the deep learning models can potentially be used to process base-wise RD data and learn to recognize specific patterns surrounding truth breakpoints**. Instead of predicting breakpoint positions directly, we formulate it as a segmentation task and infer the breakpoint positions based on the result of segmentation.

Formally, given base-wise RD vector *X* = {*d*_1_, *d*_2_, …, *d_l_*} in the screening window of *l*-bp length, we aim to find the corresponding segmentation mask vector *Y* = {*m*_1_, *m*_2_, …, *m_l_*}, in which *m_i_* ∈ {0, 1} indicates whether local coordinate *i* belongs to any SV. We propose to apply UNet [11] for learning the mapping from *X* to *Y*. The UNet is a deep neural network featured with its U-shape architecture, it integrates advantage features from the convolutional neural network (CNN) and auto-encoder (AE). For image segmentation tasks, especially medical images, it achieves the-state-of-the-art performance [12]. The structure of the UNet is described in Figure 2. It is an encode-and-decode structure that can be divided into left encoding and right decoding modules. The left-U denoises base-wise RD signals and extract features through repeatedly applying module blocks consist of a convolutional layer followed by a rectified linear unit (ReLU), batch normalization (BN). A max-pooling operation is applied in every two module blocks. The right-U expands the down-sampled feature map by up-convolution, and concatenates the correspondingly feature map from the same layer in the left-U part. Each concatenated feature map is followed by two additional convolutions with ReLU and BN. At the output layer, 1×1 convolution is applied with the *Sigmoid* function to predict mask vectors. Compared with a typical CNN model, there is no full connection layer in the UNet. Concatenating tensor in the down-sampling with the corresponding tensors in the up-sampling can help to avoid gradient vanish and keep the position features.

**Fig. 2:**
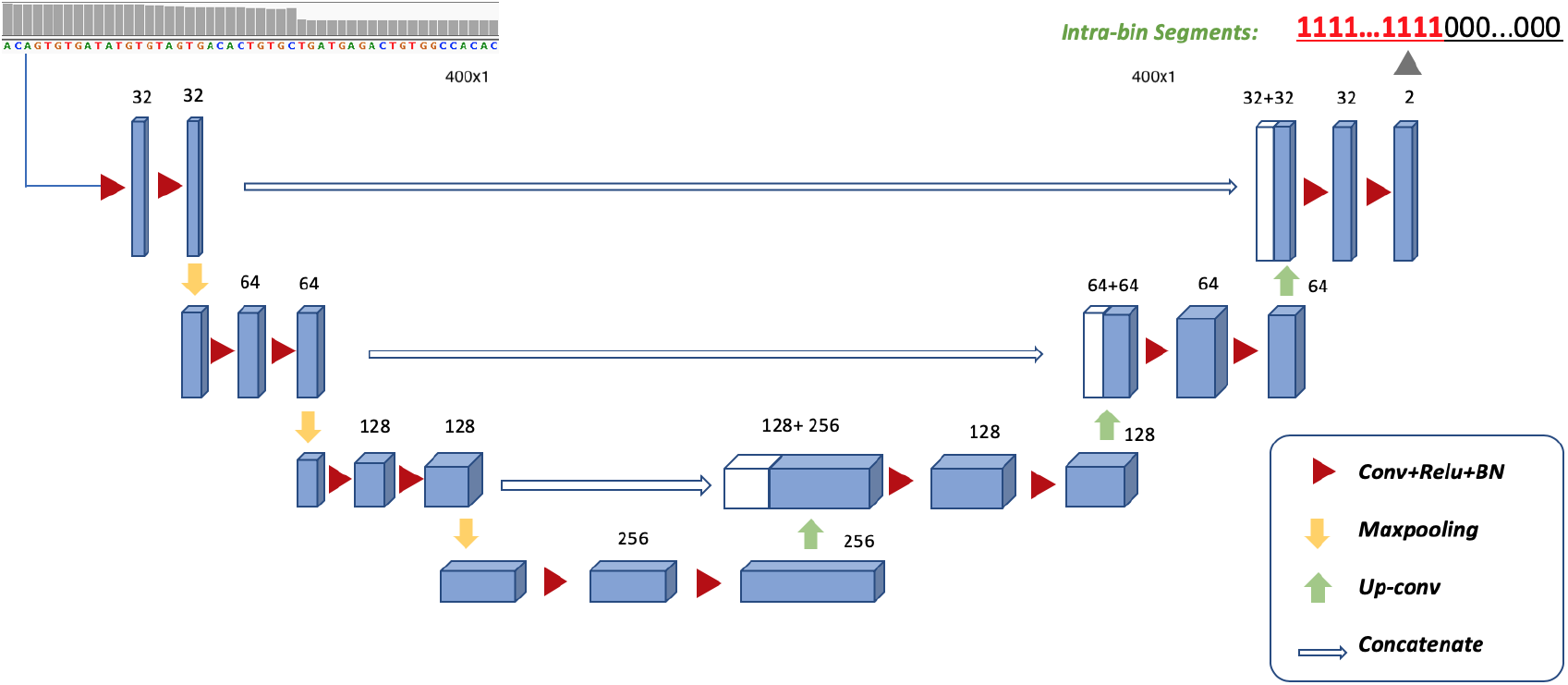
The UNet deep segmentation model used for masking SV related coordinates.

To make the prediction of breakpoint from segment result more intuitive and interpretable, we let the output of the UNet having the same length as the input. We use padding and select proper max-pooling size to avoid the non-divisible number of hidden units. The SV segment label for each position is given according to a criterion of whether the sigmoid output is larger than 0.5. If it is larger than 0.5, label “1” is assigned indicating the position belongs to an SV. Or else, label “0” is assigned for the position.

### 2.3 Training UNet for SV segmentation

To train the UNet model, we make use of known SVs and generate base-wise RD vector surrounding breakpoints of the SVs as positive samples. All-zero RD vectors are filtered out. For RD-based SV caller, SV types of deletion (DEL) and duplication (DUP) can be predicted. As the candidate breakpoint is not always in the centric position of the screening window, we take a random shift to each breakpoint as the start position of the screening window. The random shift is sampled in the range of [10, *l*−10], that a region with too small SV portion (nucleotides belongs to an SV is less than 10 bp) is treated too difficult to predict with only RD information, and excluded from the training set. Meanwhile, we generate the same amount of randomly selected RD vectors from background non-SV regions as negative samples.

We use Dice similarity coefficient (DSC) as the loss function, and Adam [13] as the optimizer to train the UNet model.

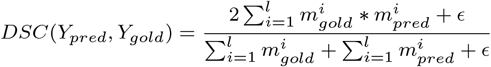

Where *s_gold_* and *s_pred_* are the gold and predicted SV label masks, and *ϵ* is a tolerance to avoid division by zero. We apply early stopping with minimum of 10 epochs and maximum of 100 epochs for the training process.

### 2.4 Apply UNet model for SV enhancement

We apply the enhancement pipeline in the following two scenarios. First, in-sample application. For a sample, we assume a small amount of SVs is validated for the sample. Those known SVs are used to train the UNet model. Then, the trained UNet model is applied to refine the rest of the non-validated SVs. Second, cross-sample application. If samples are sequenced in the same platform using similar sequencing settings, we can train the UNet model on the sample that is comprehensively investigated (e.g., NA12878) and do the enhancement for the others.

We use Algorithm 1 to infer the breakpoint position from segmentation results. As the breakpoint enhancement is performed independently for each boundary of an SV, we take a further check for the enhanced SVs, whether their SV size is less than 50 bp. If the enhancement makes an SV short, we keep the original SV boundary.

#### Algorithm 1: breakpoint enhancement

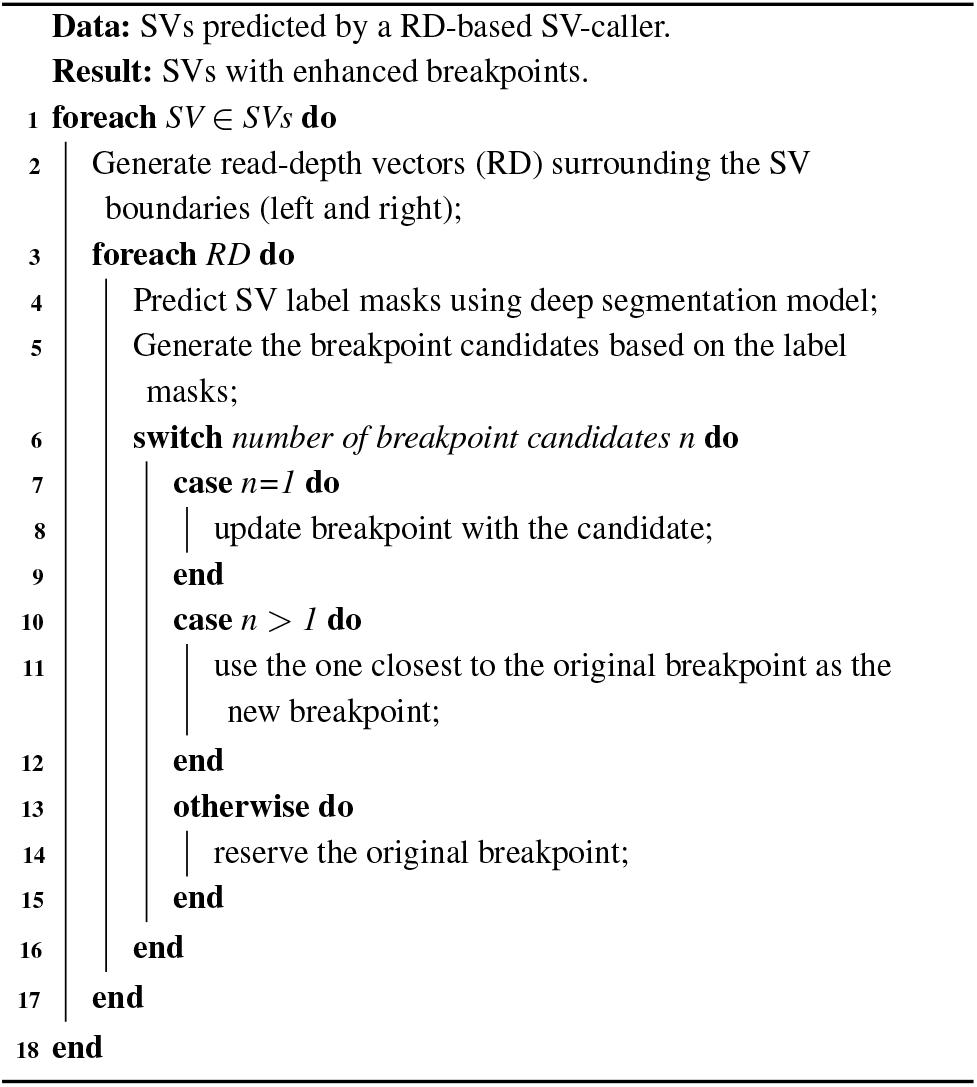

## 3 Results

### 3.1 Evaluation metrics

We evaluated the proposed method from two aspects: model and application level. On the model level, methods are evaluated by segmentation and classification task. We compared the UNet with two widely used models CNN and SVM. Typically, an SVM is used as a classifier [14], while less used as a segmentation model, Here, we only used the SVM for classification. For the segmentation models, UNet and CNN, we first segmented RD signals in the screening window and did the classification based on the segmentation results. If there is any SV mask inside of the screening window, it is classified as a breakpoint containing region, and vice versa. The DSC is used for evaluating segmentation performance. For classification, we measured with AUC, FDR, Precision, and Recall.

**Table 1.** 5-fold cross-validation results on the simulation data. Averaged result of 5-repeat runs are reported to reduce the affect of the randomness of GPU training. (Result of each run is shown in Additional File Table S5.)

| Model | Train-amount | Segmentation |  | Classification |  |  |  |
| --- | --- | --- | --- | --- | --- | --- | --- |
|  |  | DSC-ALL | DSC-BK | AUC | Precision | Recall | FDR |
| SVM | 80% | / |  | 0.8661 | 0.9102 | 0.8133 | 0.0898 |
|  | 20% | / |  | 0.8576 | 0.9002 | 0.8057 | 0.0998 |
| CNN | 80% | 0.7979 | 0.827 | 0.8759 | 0.8962 | 0.8518 | 0.1038 |
|  | 20% | 0.7478 | 0.7882 | 0.8427 | 0.8562 | 0.8266 | 0.1438 |
| UNet | 80% | **0.8240** | **0.8473** | **0.894** | **0.9288** | **0.8546** | **0.0712** |
|  | 20% | 0.8030 | 0.8311 | 0.8825 | 0.9121 | 0.8486 | 0.0879 |

On the application level, we applied the enhancement model for an RD-based SV-caller and evaluated the SV and breakpoint changes before and after that. Without loss of generality, we used the commonly used CNVnator (v 0.4) [5] as an instance. To access the accuracy of SVs, we used Jaccard similarity (JS) between predicted SV and gold standard SV, which is calculated as follows:

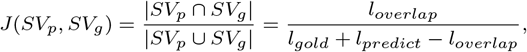

where *l_overlap_* is the length of the overlapped region between predicted and gold SV. We define a gold-overlapped SV prediction as the one with JS larger than 0.5. Compared with previous work using reciprocal overlap (RO), the length of gold SV is also considered in the Jaccard similarity. Therefore, the JS has a smaller value than that of RO. In other words, JS larger than 0.5 is a more strict metric than the RO larger than 0.5. We calculate the number of overlap SVs and SVs with precise boundary (both-boundary-match and partial-boundary-match (left or right)). **Here, the “precise” boundary is defined as within 1-bp distance to the gold.** Besides the evaluation of SVs, we also investigated the number of breakpoints with different precision. We compared the numbers of breakpoints in the pre-defined distance ranges before and after enhancement.

### 3.2 Model and related parameter settings

As in an RD-based SV-caller, the bin-size is a key parameter that affects SV prediction results. On the simulation data, we investigated several bin-sizes range from 50 bp to 1000 bp. For the enhancement model, the default length of the screening window surrounding candidate breakpoints is 400 bp. The network structure of UNet is described as in Figure 2. The convolutional neural network (CNN) uses the typical LeNet architecture. More detailed parameter information can be found in Additional File Table S1. The support vector machine (SVM) uses default settings of the Scikit-learn (v 0.22.2) package. For each WGS data, we normalized the RD vector using the mean and standard derivation of the RD in the randomly sampled background regions (non-known-SV overlapped).

### 3.3 Simulation study

We first evaluated the proposed method with simulation data. We conducted 5-fold cross-validation (CV) for evaluating model-level performance with segmentation and classification task. A typical n-fold CV uses *n* − 1 folds for training, while the rest 1-fold is used for testing. To access model performance on small-amount of data, we also evaluated the setting of training on 1-fold and testing on the rest *n* − 1 folds.

#### 3.3.1 Model-level performance

CV with the typical setting (Train-amount is 80% of SVs) was firstly investigated. Shown in Figure 1, for the segmentation task, UNet achieves better segmentation performance with higher DSC scores. For the test set containing background-regions (DSC-ALL), UNet has an absolute value of 2.61% higher than CNN. For the set only contains SV-regions (DSC-BK), UNet is around 2.03% higher than CNN. In the binary classification task, UNet achieves the best performance among the three models. UNet has higher AUCs, Precisions and Recalls while retaining lower FDRs. For deep segmentation models, UNet and CNN both have recall rate more than 85%, which are higher than SVM with the recall rate around 81.3%. But the SVM has a lower FDR than that of CNN.

#### 3.3.2 Deep segmentation models can be trained with small amount of data

In real-world applications, validated SVs are relatively few. It is worthwhile investigating training deep segmentation models with a small amount of data. For CV evaluation, we changed the typical split-setting and only used 1-fold (Train-amount:20%) for training. In the simulation data of total 19974 RD vectors, we trained models on 3989 RD vectors and tested on the rest. For those RD vectors, around half of them are generated from randomly sampled background regions that are several-distance away from the known breakpoints.

Compared with the model performance under typical CV split-setting (80% Training, 20% Testing), the performances of all three models are decreased on the smaller training data. In the segmentation task, DSC-BK of UNet and CNN have relative decreases of 1.9% and 4.7%, respectively. In the classification task, the relative AUC decrease of UNet, CNN and SVM are around 1%, 3.8%, and 1%. CNN is more sensitive to the data amount than the other two models. But the decreases are not as larger as the reduction of the training data with the amount reduced to one-quarter. This indicates that it is feasible to train such deep segmentation models with a small amount of RD-depth data. In real-world applications, we do the enhancement in the following two scenarios: One is the in-sample enhancement that trains UNet model with small amount of validated SVs, and do the breakpoint enhancement for the rest candidate SVs in the same sample. The other one is cross-sample enhancement, that trains UNet with the validated SVs from one sample, and does enhancement on another target sample. For the cross-sample case, we assume both samples are sequenced in the same sequencing platforms and settings (e.g., samples in the 1kGP), which grantees the generalization ability of the enhancement model.

#### 3.3.3 Enhancement of breakpoint resolution

We performed the in-sample enhancement for the simulation data. All models are trained on the same randomly sampled 20% of SVs. Those 20% SVs are excluded in the evaluation. CNVnator was first applied to generate initial breakpoints in bin-resolution. Then, the RD of single-nucleotide in the screening window surrounding those breakpoints are analyzed by the deep segmentation models. Based on the segmentation results, enhancement breakpoints are generated using Algorithm 1.

Table 2 demonstrates the SV results with and without enhancement. We applied CNVnator with 5 different bin-sizes range from 50 bp to 1000 bp. CNVnator using a small bin-size usually predicts more SVs, which can be observed in the column of “predicted” in Table 2. When the read-coverage of WGS data is fixed, the smaller bin-size is, the fewer reads are covered by the bin. This makes the RD information becomes noisy as the bin size decreases, especially when the bin size is less than 100 bp. For CNVnator, the number of predicted SVs is increased 21.3% to the number of 4766, when the bin-size is changed from 100 bp to 50 bp. But the number of gold-overlapped SVs does not increase in proportion to the increase of predictions. In other words, CNVnator with a smaller bin-size tends to have more false-positive predictions. Considering the additional enhancement step, we prefer to use smaller bin-size to have more SV candidates for further screening.

**Table 2.** Enhancement effect for CNVnators with different bin-sizes. The length of screening window surrounding predicted breakpoints are 400 bp. “l/r match” represents SVs with partial-boundary-match (left or right), and “l&r match” represents SVs with both-boundary-match (left and right).

| CNVnator |  | Number of specific SVs w/o Enhancement |  |  |  |  |
| --- | --- | --- | --- | --- | --- | --- |
| Bin size | predicted | enhanced | overlapped | (1) l/r match | (2) l&r match | (1)+(2) proportion |
| 50 | 4766 | / | 2237 | 182 | 4 | 8.31% |
|  |  | +CNN | 2241 | 374 | 39 | 18.43% |
|  |  | +UNet | 2256 | 725 | 866 | 70.52% |
| 100 | 3929 | / | 2227 | 101 | 3 | 4.67% |
|  |  | +CNN | 2231 | 369 | 52 | 18.87% |
|  |  | +UNet | 2239 | 704 | 887 | **71.06%** |
| 200 | 3469 | / | 2056 | 50 | 1 | 2.48% |
|  |  | +CNN | 2058 | 268 | 31 | 14.53% |
|  |  | +UNet | 2060 | 651 | 750 | 68.01% |
| 500 | 2819 | / | 1750 | 19 | 0 | 1.09% |
|  |  | +CNN | 1751 | 139 | 9 | 8.45% |
|  |  | +UNet | 1751 | 635 | 316 | 54.31% |
| 1000 | 2391 | / | 1578 | 13 | 0 | 0.82% |
|  |  | +CNN | 1580 | 74 | 1 | 4.75% |
|  |  | +UNet | 1581 | 443 | 77 | 32.89% |

In our experiment, the default screening-window length is 400 bp and a candidate breakpoint is located in the center of the window (for enhancement). For the bin-size less than 200 bp, at least one complete bin at each side of the candidate breakpoint is covered by the screening window. For the CNVnators with bin-size larger than 200 bp, the screening window may only cover a partial region of a complete bin. Therefore, the gold breakpoint may be outside the screening window for the enhancement. This explains the number of SVs with precise boundary (both-boundary-match and partial-boundary-match) reduced significantly when the bin-sizes of CNVnator are larger than 200 bp. Empirically, we make the screening window cover at least 4 bins of CNVnator (two bins at each side).

To access the enhancement effect, we investigated number changes of SVs with specific breakpoints, before and after the enhancement. From Figure 2, we can observe: (1). **The number of SVs with precise boundary (both-boundary-match and partial-boundary-match) are increased significantly**. Before enhancement, due to the bin-size limitation, SVs with precise boundary are very few. This number decreases significantly as the bin size increases. For example, CNVnator predicts at most 4 exact-boundary-match SVs with the bin-size of 50 bp. The number of partial-boundary-match SVs is less than 10%. After the enhancement, the number of exact-boundary-match SVs increases to ~887. For the CNVnator with the bin-size of 50 bp, the proportion of the precise SVs has been increased from 8.31% to 70.56% with UNet enhancement. For the enhancement using CNN, the number of precise SVs are increased to 18.43%. Those demonstrate the effectiveness and potential of using UNet for enhancing RD-based SV-callers. (2). The number of gold-overlapped SVs (JS > 0.5) is slightly increased. Compared with the CNVnator with the bin-size of 50 bp, there are 4 and 19 more gold-overlapped SVs after the enhancement using CNN and UNet, respectively. This change dues to the enhancement of new breakpoints, which makes several more SVs with Jaccard similarity larger than 0.5. (3). UNet also shows better performance than CNN that there are more partial-boundary-match SVs and significantly more exact-match-boundary SVs using the UNet enhancement.

We further investigated the enhancement effect on breakpoints. For each breakpoint, we evaluated the changes of its distance to the gold one (to-gold-distance), and counted the number of breakpoints in different to-gold-distance ranges. Figure 3 shows the number of breakpoint changes before and after enhancement with UNet and CNN. We demonstrated the change in the form of a confusing matrix and plotted in a heatmap. We split to-gold-distance in the range set of DR={[0,5), [5,10), [10,20), [20,50), [50,100), [100,200), [200, 500), [500, 1000), [1000,)}. For each element in the change matrix, *c_i,j_* represents the number of breakpoints changes from *DR_i_* to *DR_j_* after enhancement. For *c_i,i_* that breakpoints have the to-gold-distance in the same range, we ignore the unchanged breakpoints. In the figure, a cell with a large number of breakpoints is plotted in deep color. UNet enhances 2145 breakpoints with their original to-gold-distance in the range between 5 bp to 50 bp, to the breakpoints with to-gold-distance less than 5 bp, shown in the up-left corner of Figure 3(a). While the number of positive enhancement of CNN is less than that of the UNet. Note that, there exists negative enhancement that to-gold-distance increases after the enhancement. The number is smaller when compared to the number of positive enhancement. We took additional error analysis and found that most of the negative enhancements belong to the case that gold breakpoints are too far away from the initial predicted coordinate, which are also out of the screening window.

**Fig. 3:**
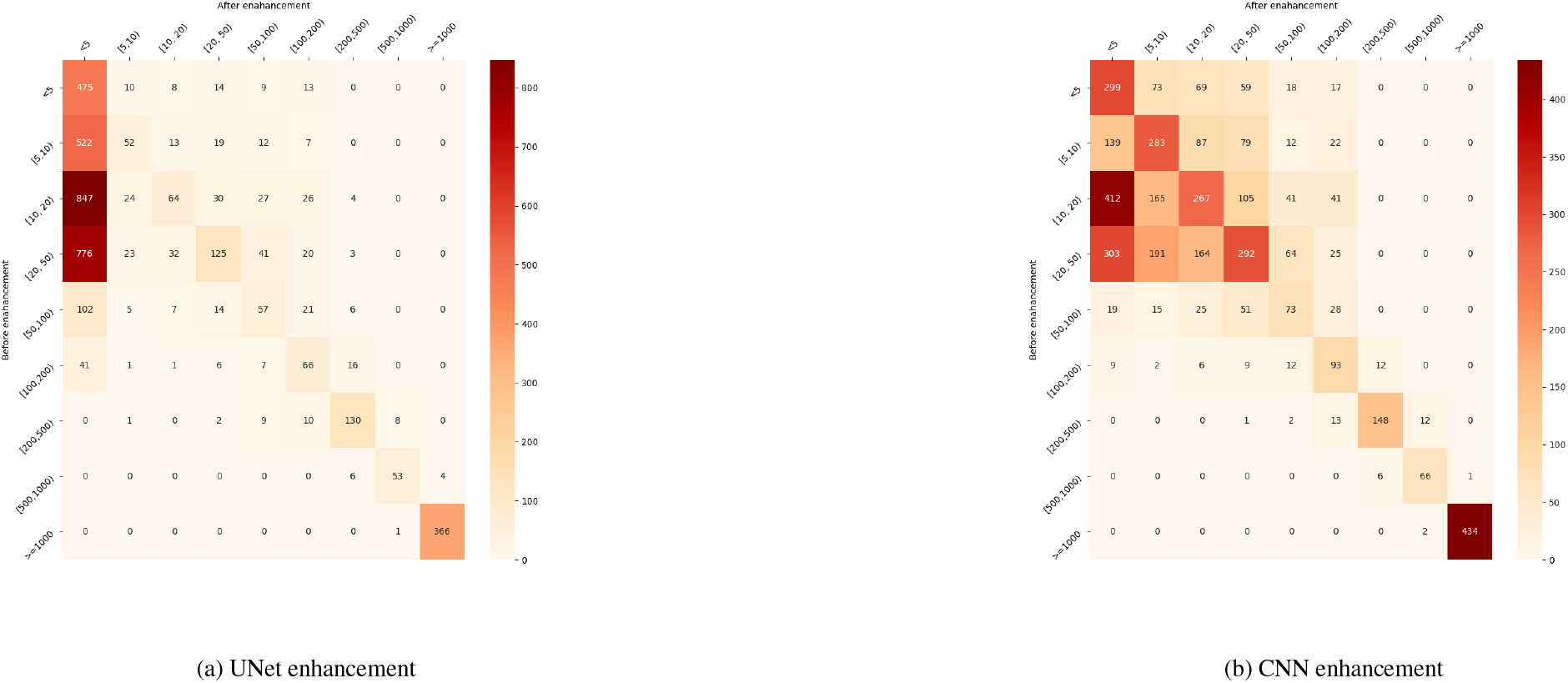
Enhancement effect by change matrix.

### 3.4 Compare enhanced RD-based SV-caller with other SV-callers

As a typical RD-based caller is not designed to predict breakpoints in single-nucleotide resolution, we compared the enhanced RD-based SV caller with Delly (v 0.8.1) [6] and Lumpy (v 0.2.13) [15], which utilize discordant and split reads to acquire single-nucleotide breakpoints. Delly predicts 3460 gold-overlapped (JS > 0.5), while Lumpy predicts 3525 gold-overlapped SVs. The number of those gold-overlapped SVs is around 1.5 times of the gold-overlapped SVs predicted by CNVnators on the same in-sample data. Around 90.82% and 92.95% of gold-overlapped SVs predicted by CNVnators are covered by Delly and Lumpy, respectively. We focused the both-boundary-match SVs predicted by different SV callers. Figure 4 shows Venn diagram of both-boundary-match SVs predicted by CNVnator (w/o enhancement) and non-RD-based SV callers. The original CNVnator without enhancement predicts only 4 both-boundary-match SVs due to the bin-size limitation. After performing RDBKE, UNet-base enhancement significantly increased the number of both-boundary-match SVs to 866. Those 866 SVs have 39.5% and 91.9% overlaps with both-boundary-match SVs predicted by Delly and Lumpy, respectively. There are 31 and 21 both-boundary-match SVs that are not overlapped with any gold-overlapped SVs of Delly and Lumpy. For the CNN enhancement, the increased number of both-boundary-match SVs (34) is much smaller than that of UNet enhancement.

**Fig. 4:**
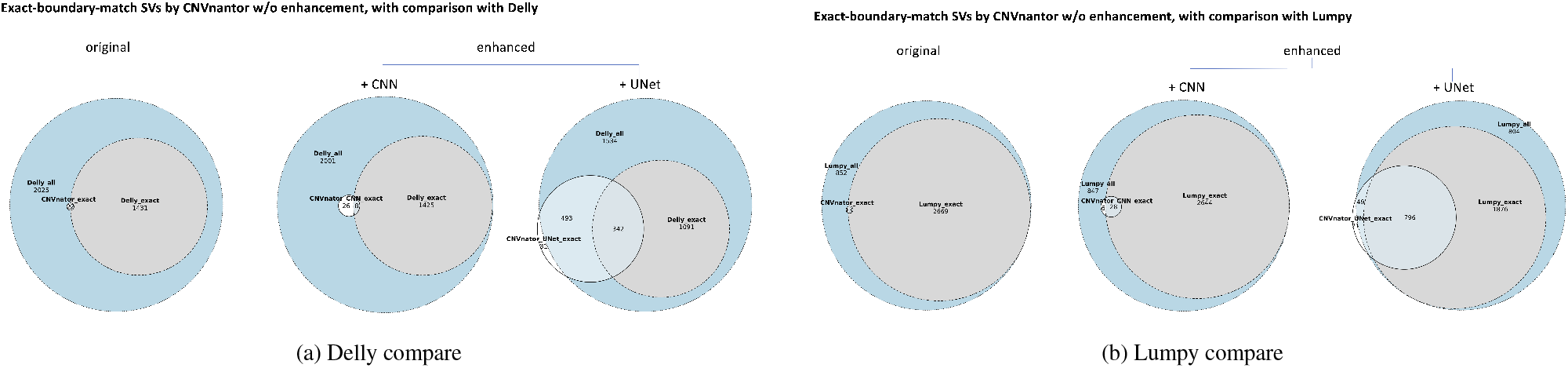
Overlaps between CNVnator (with/without enhancement) and non-RD-based SV callers (Delly and Lumpy).

### 3.5 Real data performance

We first evaluated model-level performance on the two commonly used benchmark samples: NA12878 and HG002. Shown in Figure 3, we can observe similar tendencies as in the simulation data on the segmentation and classification task. UNet gives better segmentation performance of higher DSC scores than that of CNN, which is around 2% higher on NA12878, and 2.6% ~ 3.88% higher on HG002 with different CV training split. In the classification task, UNet also achieves the best performance with all evaluation metrics than the other two models. Meanwhile, only using a one-quarter amount of the training data, the performance of UNet does not decrease significantly as that of CNN. Note that the FDR of NA12878 is higher than that of the simulation data, even though SVs of the simulation data are from the SVs of NA12878. The main reason is due to the precision of breakpoints in VCF files. In the real data, breakpoints are integrated and defined in a confidence interval that ranges from 0 bp to several hundred bp. While in the simulation data, breakpoints are precisely pre-determined. Therefore, more precise training data is important and contributes to a lower FDR of the segmentation model.

#### 3.5.1 In-sample evaluation

We conducted the in-sample evaluation on 4 real samples. For each sample, we randomly selected 20% of SVs and generated RD vectors in the non-centric screening window surrounding the boundaries of SVs for training UNet and CNN. The rest 80% SVs are used for evaluating the enhancement effect on CNVnator with 50 bp bin-size. Table 3 shows the number-changes of specific SVs and breakpoints before and after the enhancement. For the number of SVs with specific boundaries, UNet gives more precise-boundary SVs than that of CNN on all the 4 samples. For the sample of HG002, the total share of partial-boundary-match SVs and both-boundary-match SVs is around 59.72%. Besides, we can observe the number of breakpoints with to-gold-distance less than 5 bp is also increased significantly after the UNet enhancement. The mean of to-gold-distances less than 5bp are also reduced, shown in Additional File Table S3. The change-matrices like Figure 3, are shown in Additional File Figure S1. Most of the positive enhancements of UNet are for those breakpoints with the original to-gold-distance in the range between 20 bp and 100 bp. While for CNN, the enhancement is conservative that less number of breakpoints are enhanced with the to-gold-distance less than 5 bp.

**Table 3.** Model-level evaluation on the real data. 5-fold cross-validation results on NA12878 and HG002 with screening window length of 400 bp. Averaged result of 5-repeat runs are reported to reduce the affect of the randomness of GPU training.

| Model | Train-amount | NA12878 |  |  |  |  |  | HG002 |  |  |  |  |  |
| --- | --- | --- | --- | --- | --- | --- | --- | --- | --- | --- | --- | --- | --- |
|  |  | Segmentation |  | Classification |  |  |  | Segmentation |  | Classification |  |  |  |
|  |  | DSC-ALL | DSC-BK | AUC | Precision | Recall | FDR | DSC-ALL | DSC-BK | AUC | Precision | Recall | FDR |
| SVM | 80% | / |  | 0.838 | 0.8403 | 0.834 | 0.1597 | / |  | 0.8822 | 0.891 | 0.8709 | 0.109 |
|  | 20% | / |  | 0.8331 | 0.8351 | 0.8296 | 0.1649 | / |  | 0.8765 | 0.8885 | 0.8614 | 0.1115 |
| CNN | 80% | 0.7688 | 0.8066 | 0.8557 | 0.8595 | 0.8505 | 0.1405 | 0.8024 | 0.8277 | 0.8947 | 0.8880 | 0.9035 | 0.1120 |
|  | 20% | 0.7466 | 0.7862 | 0.8405 | 0.8575 | 0.8174 | 0.1425 | 0.7719 | 0.7987 | 0.8854 | 0.8842 | 0.8880 | 0.1158 |
| UNet | 80% | 0.7898 | 0.8257 | 0.8698 | 0.8805 | 0.8566 | 0.1183 | 0.8284 | 0.8520 | 0.9122 | 0.9163 | 0.9076 | 0.0837 |
|  | 20% | 0.7696 | 0.8098 | 0.8608 | 0.8709 | 0.8480 | 0.1291 | 0.8107 | 0.8358 | 0.9027 | 0.9087 | 0.8963 | 0.0913 |

**Table 4.**
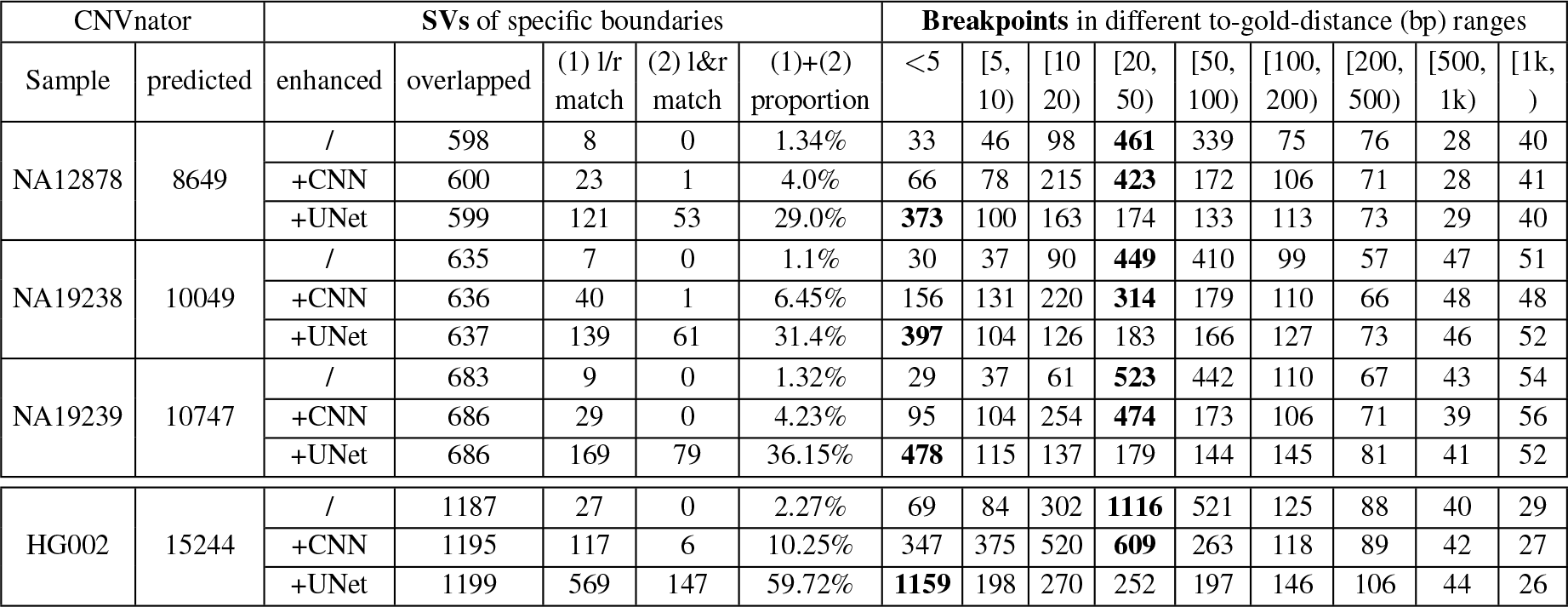
In-sample evaluation of the enhancement for CNVNator with 50 bp bin-size. The largest number of breakpoints of each SV caller is highlighted in bold font. “l/r match” represents SVs with partial-boundary-match (left or right), and “l&r match” represents SVs with both-boundary-match (left and right).

#### 3.5.2 Cross-sample evaluation

We evaluated cross-sample enhancement for the WGS data from 1kGP. The key preliminary of applying cross-sample enhancement is the deep segmentation model can be generalized across samples. Therefore, for the cross-sample enhancement, we assume those samples are sequenced in the same platform with a similar sequencing setting (e.g., read-length). In other words, we assume both samples share a similar whole-genome read coverage. This is not a rare case for WGS data generated in mainstream sequencing platforms or WGS data generated in a consortium study. We used comprehensively investigated NA12878 to train deep segmentation models, and applied for the samples of NA19238 and NA19239. Different from the in-sample evaluation that 20% SVs used in the training are not evaluated, all known SVs of the sample are evaluated in the cross-sample evaluation.

Table 5 shows the cross-sample evaluation on model-level and enhancement-level. On the model-level, for all models on the classification task, we can observe recalls are lower, while precision scores retain referring to the in-sample evaluation. UNet also achieves the best segmentation and classification performance among the three models. For SVM, the FDR has the worst score of 0.2045 and 0.1626 on NA19238 and NA19239, respectively. This indicates that for the cross-sample application, directly classification based on the RD information is not as robust as the segmentation based approach that focuses on the local RD shift patterns. On the enhancement-level, similar enhancement effect like the in-sample evaluation can be observed for the UNet model. The proportion of SVs with precise boundaries increases from around 1.3% to around 35%. The mean and standard derivation of to-gold-distance in different ranges are shown in Additional File Table S4. The performance of CNN has obvious improvement when compared with the in-sample enhancement. Breakpoint change-matrices are shown in Additional File Table S2. More breakpoints are enhanced with to-gold-distance less than 5 bp. As discussed in the previous section, CNN is more sensitive to the amount of training data. In the cross-sample evaluation, more amount of data is available for the training, which contributes to the performance improvement of CNN. But the number of SVs with precise boundaries and the number of breakpoints with to-gold-distance less than 5 bp are still less than that of UNet.

**Table 5.**
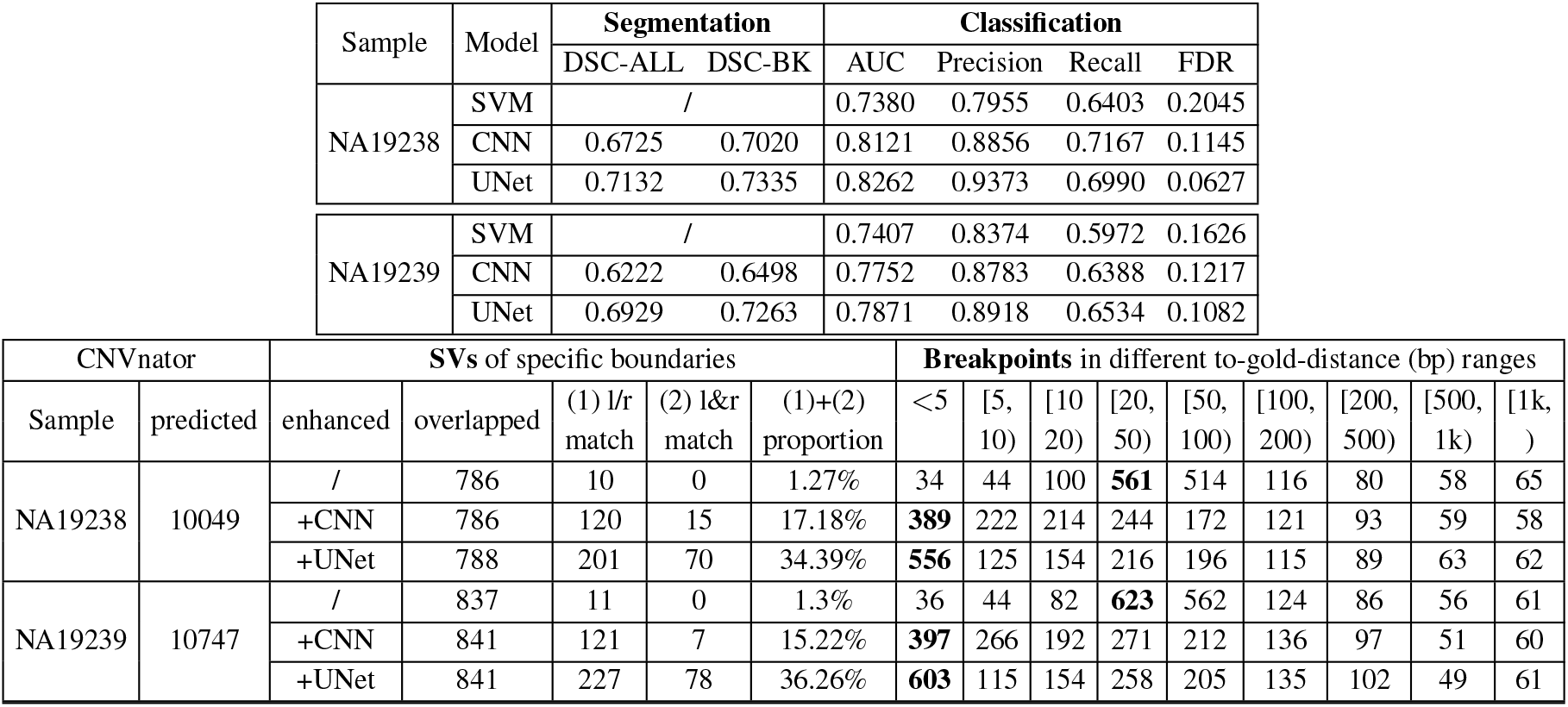
Cross-sample evaluation on real data. Deep segmentation models are trained on NA12878 and enhancement are applied for NA19238 and NA19239. “l/r match” represents SVs with partial-boundary-match (left or right), and “l&r match” represents SVs with both-boundary-match (left and right).

### 3.6 Effect of read-depth and screening window length

#### 3.6.1 Effect of read-depth

The enhancement effect is affected by the read-depth and screening window length. We empirically evaluated how read-depth affects model performance through down-sampling the NA12878 WGS data. We conducted 5-fold cross-validation with 20% train-split for NA12878 with different down-sampling rate. In Figure 5, we can observe a general trend of performance improvement as the increase of read-depth. Curves in the region of the down-sampling rate below 0.5 are relatively steeper compared with curves in the rest region. It indicates the difficulty of training a deep segmentation model for a single sample using low-depth data. Based on performance gaps shown in the figure, more than 40x are empirically preferred minimal read-depth for applying the enhancement model.

**Fig. 5:**
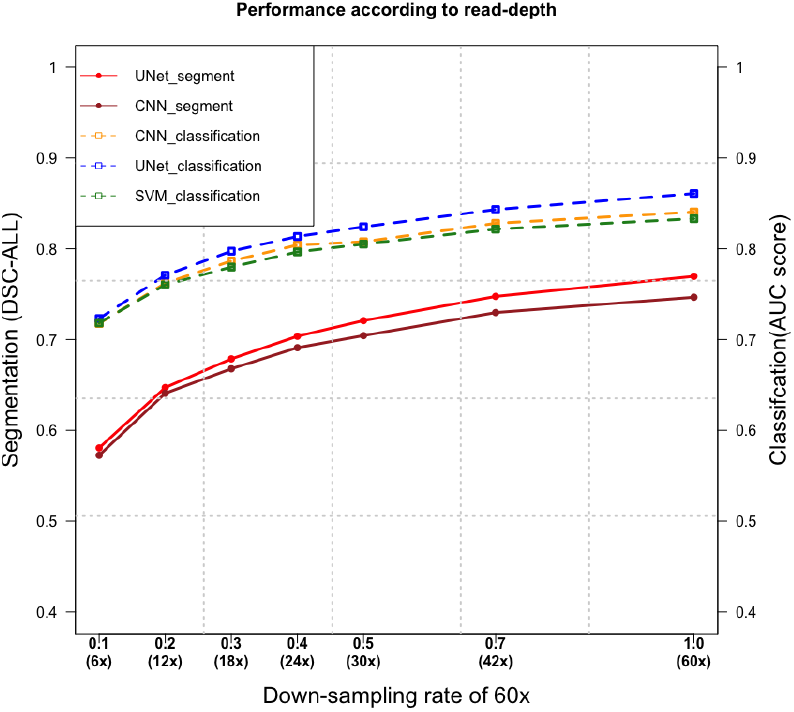
Performance of different read-depths through down-sampling 60x WGS of NA12878. The dashed curves connect AUC scores of classification, while the line curves show the DSC-ALL scores of segmentation.

#### 3.6.2 Effect of screening window length

To evaluate the effect of different screening window lengths, we used NA12878 to do in-sample enhancement for 50 bp bin-size CNVnator. The screening window length ∈ {100*bp*, 200*bp*, 400*bp*, 800*bp*, 1000*bp*} are evaluated. Shown in Figure 6, on the model-level performance, different screening window length does not make drastic performance changes like the effect of the read-depth. On the classification task, we can observe the performances of SVM and CNN slightly decrease as the length of the screening window increases, while the AUC curve of UNet starts to diverge after the length of 200 bp and has a converged trend with higher AUC scores than that of SVM and CNN. The longer a screening window is, the more RD information is included. SVM and CNN seem to be more affected by more noisy non-SV related RD signals, while UNet generalizes well for the longer screen window lengths. On the segmentation task, as the increase of screening window length, the number of evaluated label masks is also increased. The overall DSC scores are increased accordingly. UNet performs better than CNN, especially with the screening window length larger than 400 bp.

**Fig. 6:**
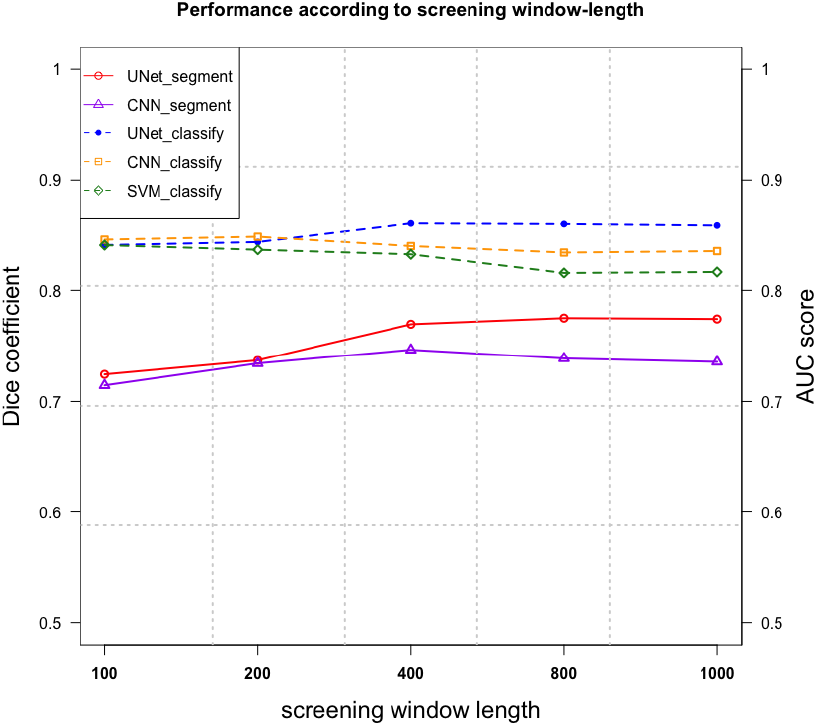
Performance of different screening window length for NA12878 data. The dashed curves connect AUC scores of classification, while the line curves show the DSC-ALL scores of segmentation.

## 4 Discussion

In this paper, we proposed RDBKE, a general enhancement approach to increase the breakpoint resolution for read-depth based SV callers. The key component of RDBKE is using the UNet model to segment regions surrounding candidate breakpoints. Previous RD-based SV callers require smoothed RD signals by bins, which limits the breakpoint resolution. Here, we used UNet to process base-wise RD signals directly and derived single-nucleotide resolution breakpoint predictions. Although the single-base RD is very noisy for the short-read sequencing data, utilization of a deep learning model with a proper neural network structure can be used to process such noisy data. Besides the convolutional modules in hierarchical levels, the encoding-decoding architecture and skip-connection also contribute to this functionality. Besides, the UNet model can be trained with a small amount of training data, which makes the in-sample application feasible. The enhancement pipeline can be also applied for cross-sample with more training data under the condition that both samples are sequenced in the same platform with a similar sequencing setting.

Recent SV benchmark studies [9, 16] show that no single SV caller can accurately and sensitively detect all types and all sizes of SVs. To further improve SV prediction, an SV callset is commonly generated through clustering SV predictions of different SV callers. For those benchmark SV callsets, multiple sequencing technologies are also applied for the same sample to derive high-confidence results. To filter and merge SV results, different computational methods are developed [17, 14]. For example, SVclassify [14] uses 1-class SVM to cluster and classify whether candidate SVs have abnormal annotations different from most of the genome. Here, we focused on the RD-based SV caller. Instead of developing a new RD-based SV caller, we proposed to use deep learning models for enhancing the existing RD-based SV caller. Related but different from existed machine learning applications for SV detection as the classification task, we took a different modeling approach as the segmenting, which can provide a finer granularity analysis for putative regions.

Comparing with other types of read signals, RD information is more common. Although the split-read algorithm can accurately predict base-wise breakpoints, there is a limited number of SVs that are covered by plenty of split reads [18]. Pedersen et al., [19] proposed to integrated read-depth information to analyze putative events generated through the clustering of discordant and split-read algorithms. In their methods, they compared the median depth in the event to the median depth from the 1000 bases on either side, which is further used to refine predictions of split-read and paired-end based methods. This method uses bin-based RD information.

We took further error analysis for the output of the segmentation models on the simulation data. For those regions containing drastic RD changes, it is relatively easy for both UNet and CNN to label SV-related coordinates. UNet tends to make more consecutive label masks than that of CNN, especially near the candidate boundaries. For those regions with less drastic RD change, it is still possible for both deep segmentation models to make almost correct segments, shown in Additional File Figure S3 (a). Both segmentation models show cases that work for small size SVs (Figure S3 (b)). Besides the gold breakpoints that are outside the screening window, two possible issues are related to the observed errors. One is the RD signal is less informative to make reliable segmentation. The other is inconsistent correlation between the breakpoints and RD signals, shown in Figure S3 (c). Although the segmentation is not perfect, our experiments demonstrate that base-wise RD information can still be used to learn specific signal patterns surrounding breakpoints. The performance will improve when high-depth short-read sequencing data and high-quality training data become more and more available. To alleviate the effect of wrong segments, the original SVs can be retained with the enhancement for further clustering-based analysis.

In this work, we only used general RD information to enhance the breakpoint resolution of SVs. The proposed deep learning framework also has the flexibility of incorporating other heterogeneous features as input. For example, we may incorporate sequence-related information and read-depth of specific reads, such as split-reads and paired-end read. Recently, deep variant [10] uses pileup images of putative regions (100 bp) and applied CNN to classify genotypes for detecting SNP and small indel variants. Although pileup images representation contain more information, they are noisier than 1-dimensional RD data, especially when putative regions become larger in the SV detection task. It is worthwhile being further explored.

## 5 Conclusions

In this paper, we present RDBKE for enhancing breakpoints with the single-nucleotide resolution for a general RD-based SV caller. The enhancement is achieved through learning base-wise RD patterns surrounding known breakpoints with the deep segmentation model UNet. We show that UNet can be trained with a small amount of data and apply for breakpoint enhancement in-sample and cross-sample. On both simulation and real data, RDBKE with UNet significantly increases the number of SVs with more precise breakpoints.

## Additional File

Additional File1: Table S1-S5, Figure S1-S3.

