## Additional File Table S1-S5, Figure S1-S3 for "Enhancing breakpoint resolution with deep segmentation model: a general refinement method for read-depth based structural variant callers": Supplements.pdf

### Table S1. (a). Detailed neural network structural of UNet

| Layer (type) | Output Shape | Param # | Connected to |
|---|---|---|---|
| input (InputLayer) | (None, 400, 1) | 0 | |
| conv1d_1 (Conv1D) | (None, 400, 32) | 384 | input[0][0] |
| batch_normalization_1 (BatchNor | (None, 400, 32) | 128 | conv1d_1[0][0] |
| conv1d_2 (Conv1D) | (None, 400, 32) | 11296 | batch_normalization_1[0][0] |
| batch_normalization_2 (BatchNor | (None, 400, 32) | 128 | conv1d_2[0][0] |
| max_pooling1d_1 (MaxPooling1D) | (None, 80, 32) | 0 | batch_normalization_2[0][0] |
| conv1d_3 (Conv1D) | (None, 80, 64) | 22592 | max_pooling1d_1[0][0] |
| batch_normalization_3 (BatchNor | (None, 80, 64) | 256 | conv1d_3[0][0] |
| conv1d_4 (Conv1D) | (None, 80, 64) | 45120 | batch_normalization_3[0][0] |
| batch_normalization_4 (BatchNor | (None, 80, 64) | 256 | conv1d_4[0][0] |
| max_pooling1d_2 (MaxPooling1D) | (None, 16, 64) | 0 | batch_normalization_4[0][0] |
| conv1d_5 (Conv1D) | (None, 16, 128) | 90240 | max_pooling1d_2[0][0] |
| batch_normalization_5 (BatchNor | (None, 16, 128) | 512 | conv1d_5[0][0] |
| conv1d_6 (Conv1D) | (None, 16, 128) | 180352 | batch_normalization_5[0][0] |
| batch_normalization_6 (BatchNor | (None, 16, 128) | 512 | conv1d_6[0][0] |
| max_pooling1d_3 (MaxPooling1D) | (None, 8, 128) | 0 | batch_normalization_6[0][0] |
| conv1d_7 (Conv1D) | (None, 8, 256) | 360704 | max_pooling1d_3[0][0] |
| batch_normalization_7 (BatchNor | (None, 8, 256) | 1024 | conv1d_7[0][0] |
| conv1d_8 (Conv1D) | (None, 8, 256) | 721152 | batch_normalization_7[0][0] |
| batch_normalization_8 (BatchNor | (None, 8, 256) | 1024 | conv1d_8[0][0] |
| up_sampling1d_1 (UpSampling1D) | (None, 16, 256) | 0 | batch_normalization_8[0][0] |
| concatenate_1 (Concatenate) | (None, 16, 384) | 0 | batch_normalization_6[0][0]<br>up_sampling1d_1[0][0] |
| conv1d_9 (Conv1D) | (None, 16, 128) | 540800 | concatenate_1[0][0] |
| batch_normalization_9 (BatchNor | (None, 16, 128) | 512 | conv1d_9[0][0] |
| conv1d_10 (Conv1D) | (None, 16, 128) | 180352 | batch_normalization_9[0][0] |
| batch_normalization_10 (BatchNo | (None, 16, 128) | 512 | conv1d_10[0][0] |
| up_sampling1d_2 (UpSampling1D) | (None, 80, 128) | 0 | batch_normalization_10[0][0] |
| concatenate_2 (Concatenate) | (None, 80, 192) | 0 | batch_normalization_4[0][0]<br>up_sampling1d_2[0][0] |
| conv1d_11 (Conv1D) | (None, 80, 64) | 135232 | concatenate_2[0][0] |
| batch_normalization_11 (BatchNo | (None, 80, 64) | 256 | conv1d_11[0][0] |
| conv1d_12 (Conv1D) | (None, 80, 64) | 45120 | batch_normalization_11[0][0] |
| batch_normalization_12 (BatchNo | (None, 80, 64) | 256 | conv1d_12[0][0] |
| up_sampling1d_3 (UpSampling1D) | (None, 400, 64) | 0 | batch_normalization_12[0][0] |
| concatenate_3 (Concatenate) | (None, 400, 96) | 0 | batch_normalization_2[0][0]<br>up_sampling1d_3[0][0] |
| conv1d_13 (Conv1D) | (None, 400, 32) | 33824 | concatenate_3[0][0] |
| batch_normalization_13 (BatchNo | (None, 400, 32) | 128 | conv1d_13[0][0] |
| conv1d_14 (Conv1D) | (None, 400, 32) | 11296 | batch_normalization_13[0][0] |
| batch_normalization_14 (BatchNo | (None, 400, 32) | 128 | conv1d_14[0][0] |
| conv1d_15 (Conv1D) | (None, 400, 2) | 706 | batch_normalization_14[0][0] |
| batch_normalization_15 (BatchNo | (None, 400, 2) | 8 | conv1d_15[0][0] |
| conv1d_16 (Conv1D) | (None, 400, 1) | 3 | batch_normalization_15[0][0] |

Total params: 2,384,813
Trainable params: 2,381,993
Non-trainable params: 2,820

### Table S1. (b). Detailed neural network structural of CNN

| Layer (type) | Output Shape | Param # |
|---|---|---|
| conv1d_1 (Conv1D) | (None, 390, 64) | 768 |
| max_pooling1d_1 (MaxPooling1 | (None, 78, 64) | 0 |
| conv1d_2 (Conv1D) | (None, 72, 128) | 57472 |
| max_pooling1d_2 (MaxPooling1 | (None, 14, 128) | 0 |
| flatten_1 (Flatten) | (None, 1792) | 0 |
| dense_1 (Dense) | (None, 256) | 459008 |
| dropout_1 (Dropout) | (None, 256) | 0 |
| dense_2 (Dense) | (None, 400) | 102800 |

Total params: 620,048
Trainable params: 620,048
Non-trainable params: 0

### Table S2. (a).VCF files used in the evaluation

| Samples | VCF files |
|---|---|
| Simulation | https://github.com/stat-lab/EvalSVcallers/blob/master/Ref_SV/Sim-A.SV.vcf |
| NA12878, NA19238, NA19239 | ftp://ftp.1000genomes.ebi.ac.uk/vol1/ftp/phase3/integrated_sv_map/ALL.wgs.mergedSV.v8.20130502.svs.genotypes.vcf.gz |
| HG002 | ftp://ftp-trace.ncbi.nlm.nih.gov/giab/ftp/data/AshkenazimTrio/analysis/NIST_SVs_Integration_v0.6/HG002_SVs_Tier1_v0.6.vcf.gz |

**(b). BAM files**

| Samples | BAM files |
|---|---|
| NA12878 | ftp://ftp.1000genomes.ebi.ac.uk/vol1/ftp/data/NA12878/high_coverage_alignment/NA12878.mapped.ILLUMINA.bwa.CEU.high_coverage_pcr_free.20130906.bam |
| NA19238 | ftp://ftp.1000genomes.ebi.ac.uk/vol1/ftp/data/NA12878/high_coverage_alignment/NA19238.mapped.ILLUMINA.bwa.YRI.high_coverage_pcr_free.20130924.bam |
| NA19239 | ftp://ftp.1000genomes.ebi.ac.uk/vol1/ftp/data/NA12878/high_coverage_alignment/NA19239.mapped.ILLUMINA.bwa.YRI.high_coverage_pcr_free.20130924.bam |
| HG002 | ftp://ftp-trace.ncbi.nlm.nih.gov/giab/ftp/data/AshkenazimTrio/HG002_NA24385_son/NIST_HiSeq_HG002_Homogeneity-10953946/NHGRI_Illumina300X_AJtrio_novoalign_bams/HG002.hs37d5.60X.1.bam. |

**Figure S1. Breakpoint change matrix of in-sample enhancement.**

**NA12878**

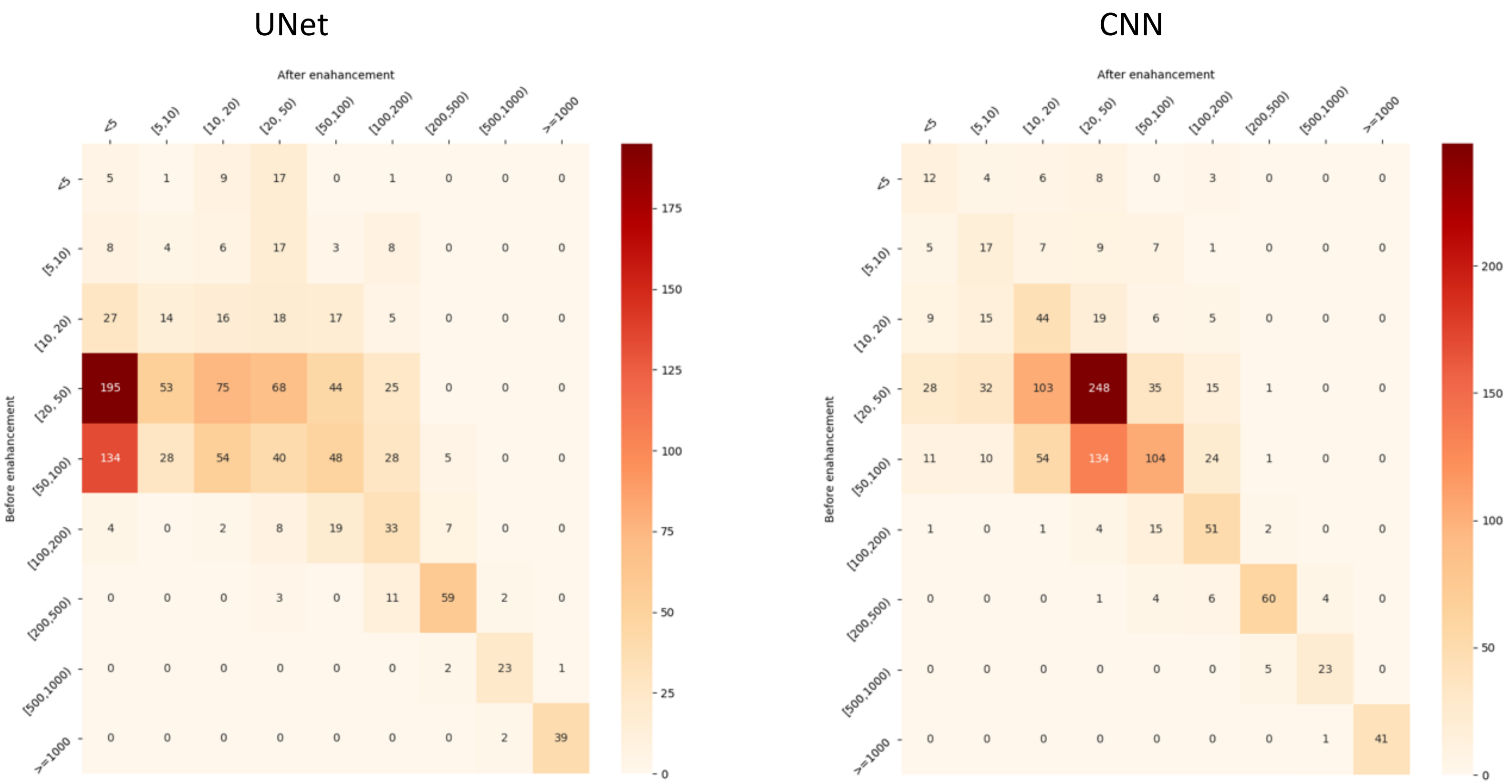

**NA19238**

UNet

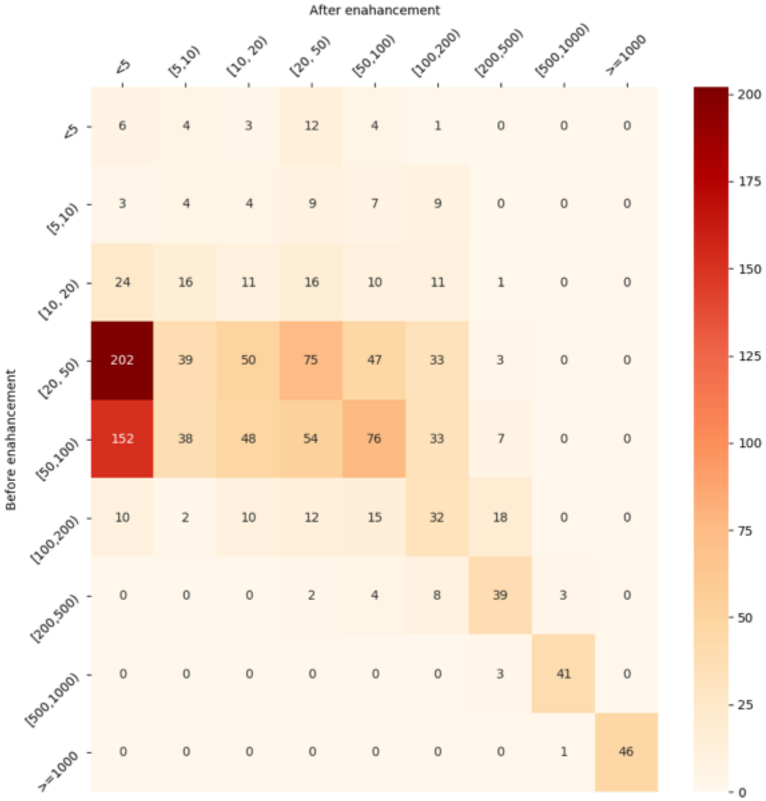

CNN

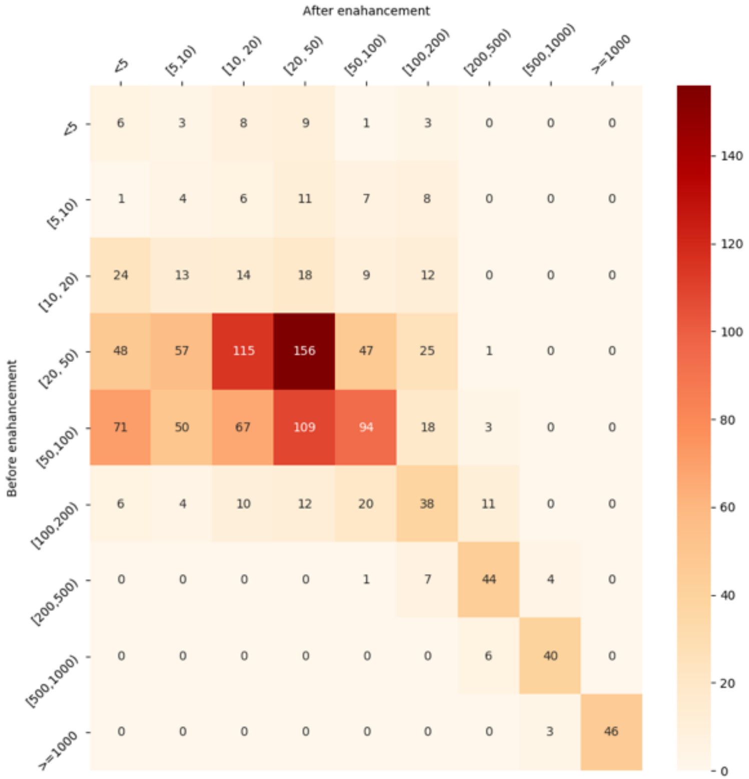

**NA19239**

UNet

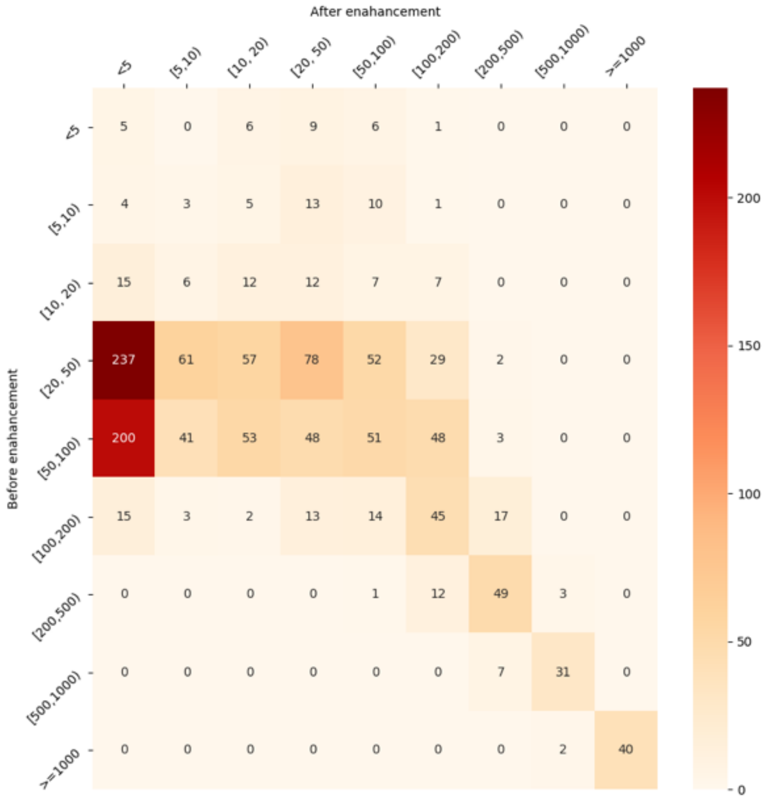

CNN

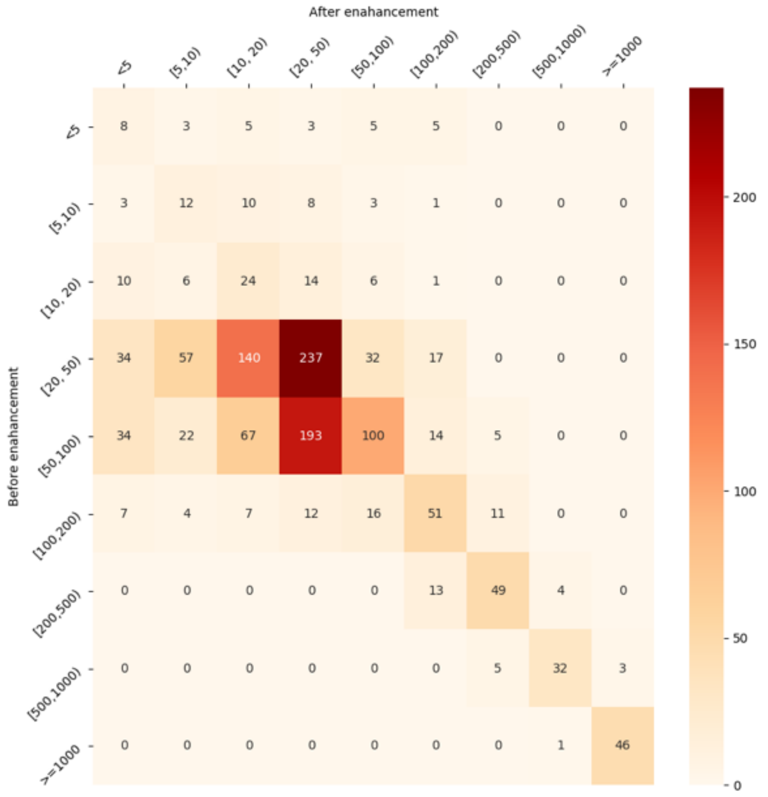

## HG002

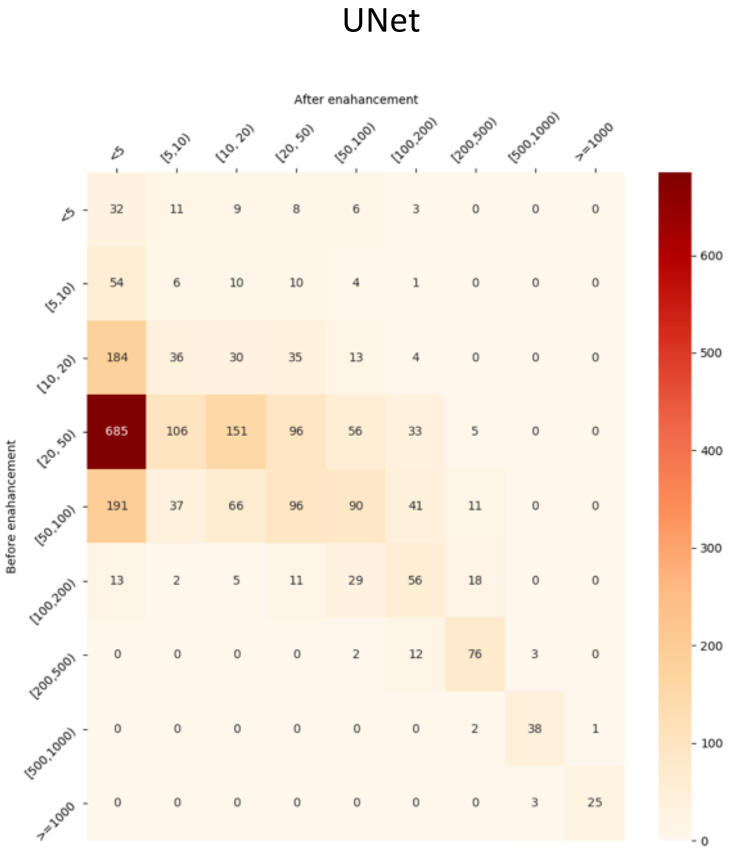

UNet
After enahancement
Before enahancement

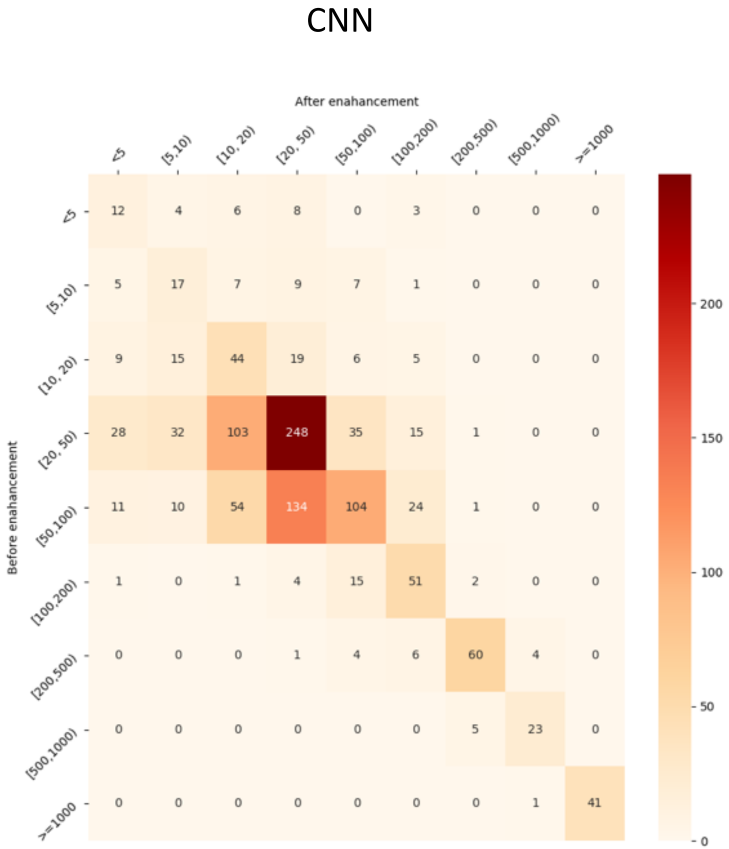

CNN
After enahancement
Before enahancement

**Figure S2. Breakpoint change matrix of cross-sample enhancement.**

**<u>NA19238</u>**

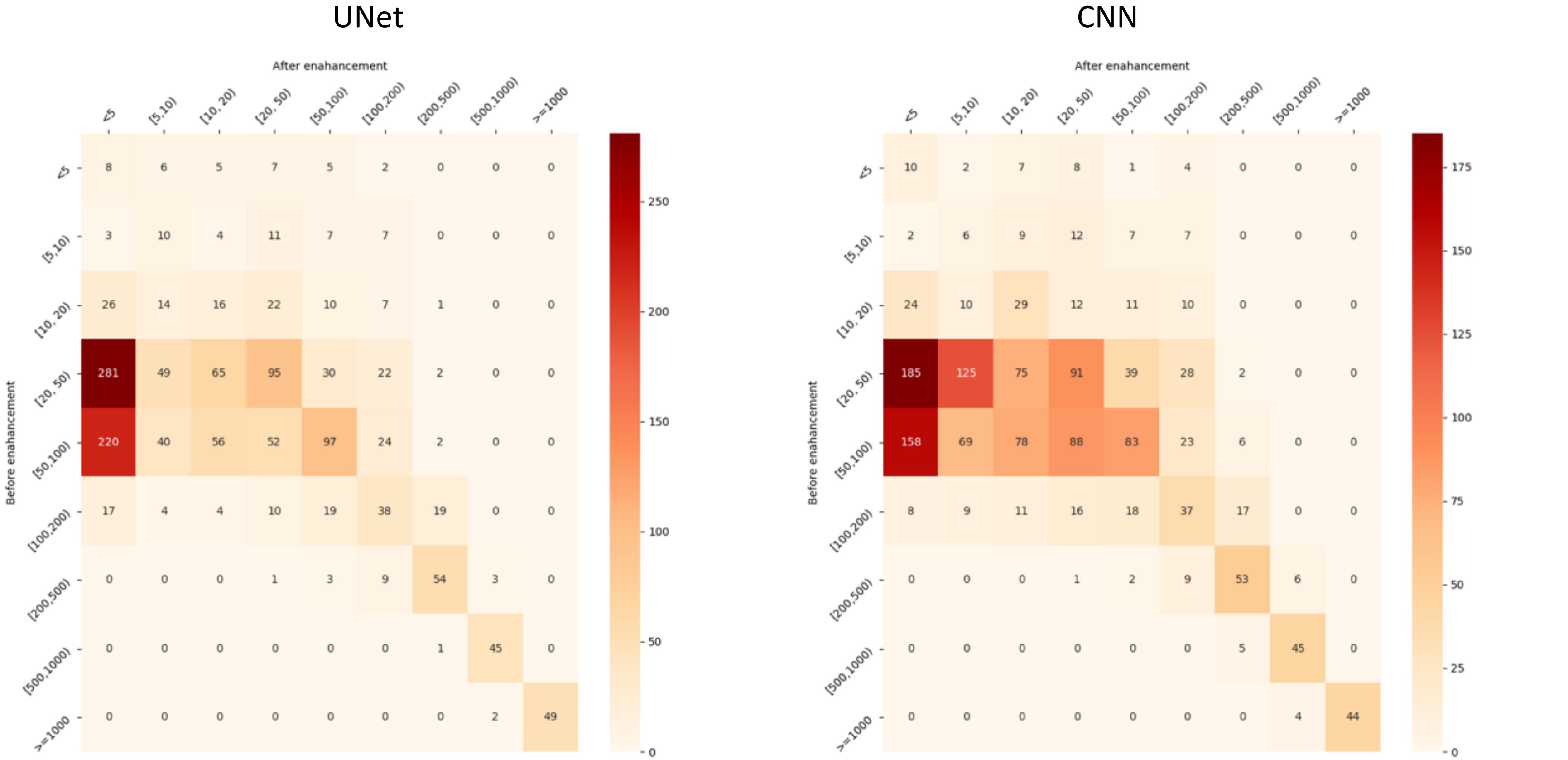

**NA19239**

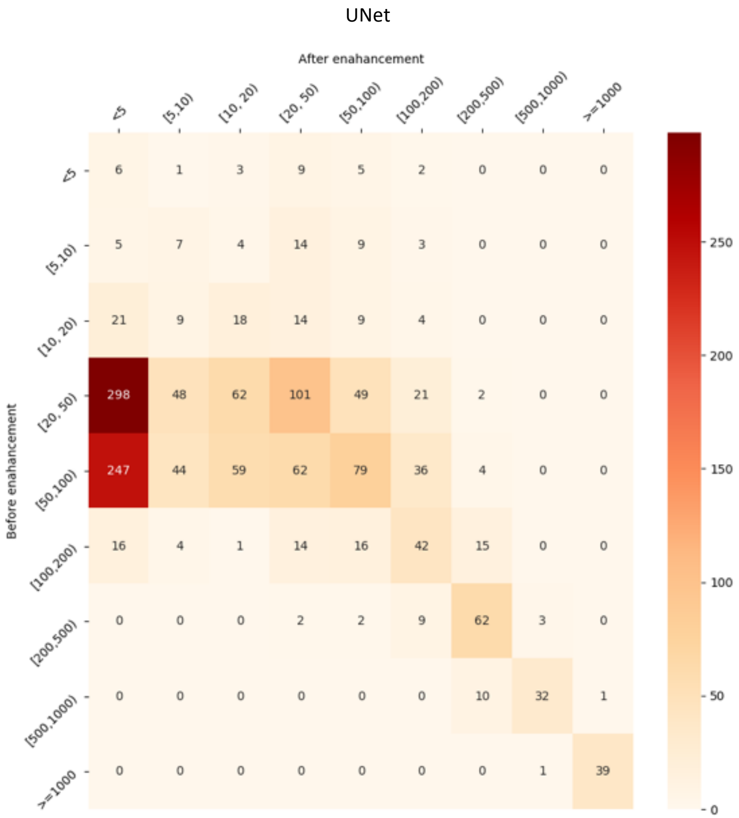
UNet
After enahancement
<5 [5,10) [10, 20) [20, 50) [50,100) [100,200) [200,500) [500,1000) >=1000
Before enahancement
<5 6 1 3 9 5 2 0 0 0
[5,10) 5 7 4 14 9 3 0 0 0
[10, 20) 21 9 18 14 9 4 0 0 0
[20, 50) 298 48 62 101 49 21 2 0 0
[50,100) 247 44 59 62 79 36 4 0 0
[100,200) 16 4 1 14 16 42 15 0 0
[200,500) 0 0 0 2 2 9 62 3 0
[500,1000) 0 0 0 0 0 0 10 32 1
>=1000 0 0 0 0 0 0 0 1 39
250
200
150
100
50
0

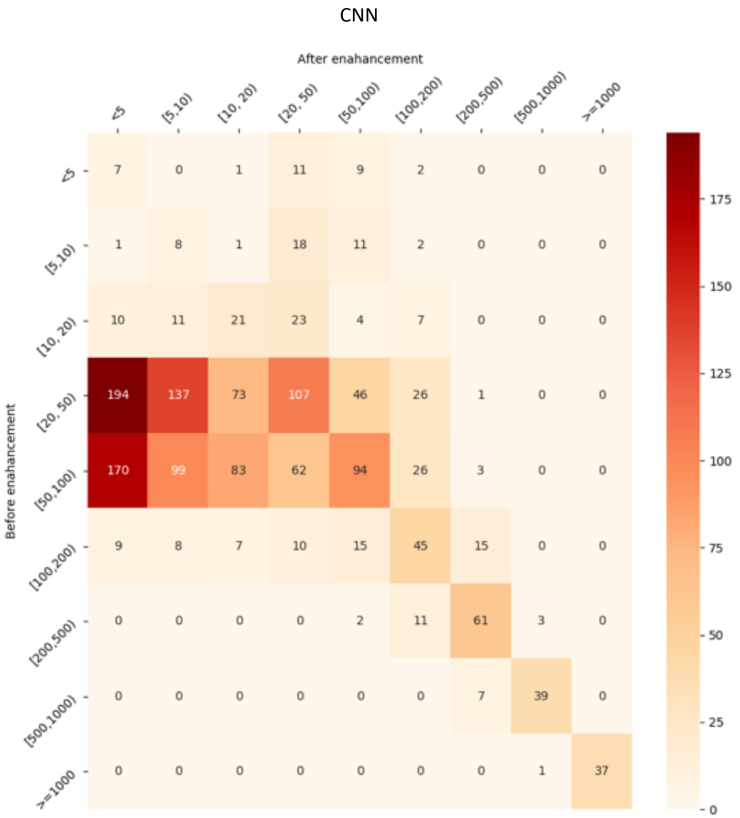
CNN
After enahancement
<5 [5,10) [10, 20) [20, 50) [50,100) [100,200) [200,500) [500,1000) >=1000
Before enahancement
<5 7 0 1 11 9 2 0 0 0
[5,10) 1 8 1 18 11 2 0 0 0
[10, 20) 10 11 21 23 4 7 0 0 0
[20, 50) 194 137 73 107 46 26 1 0 0
[50,100) 170 99 83 62 94 26 3 0 0
[100,200) 9 8 7 10 15 45 15 0 0
[200,500) 0 0 0 0 2 11 61 3 0
[500,1000) 0 0 0 0 0 0 7 39 0
>=1000 0 0 0 0 0 0 0 1 37
175
150
125
100
75
50
25
0

### Figure S3. Segmentation examples on Simulation the data. Dash line indicates the position of the gold breakpoint. Coordinates in red are the SV related masks.

#### (a). Examples of positive enhancement by UNet

**obvious case**

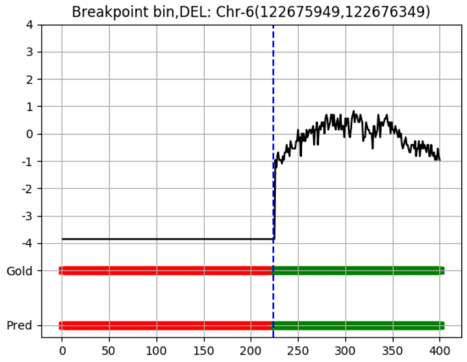

**less obvious case**

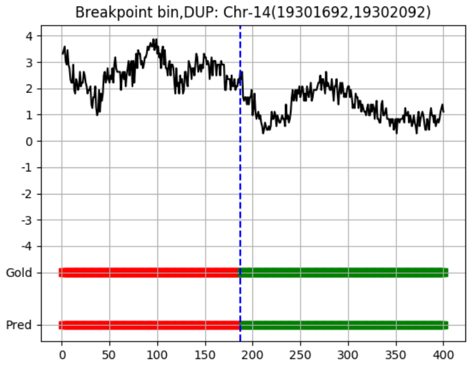

#### (b). small SVs totally inside the screening window

#### Unet

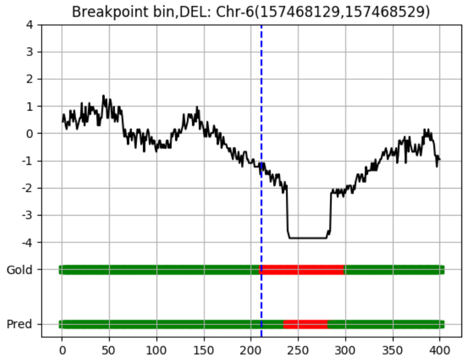

**Left**

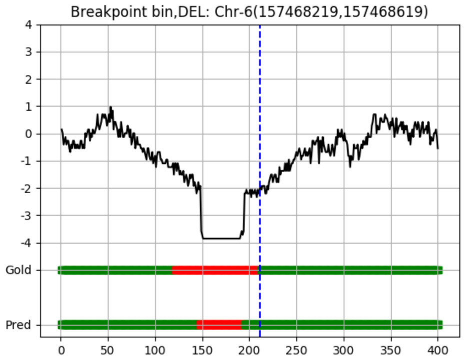

**Right**

**CNN**

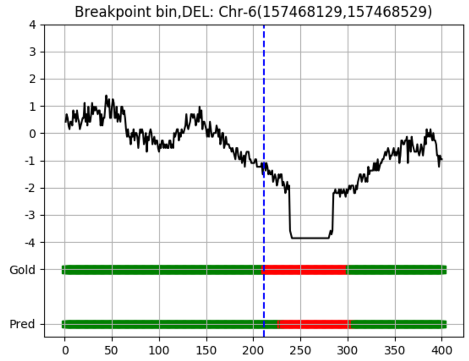

**Left**

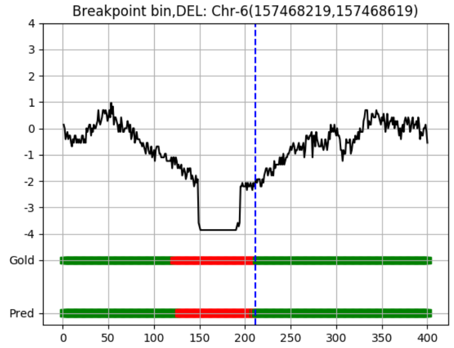

**Right**

### (c). Negative segmentation of UNet

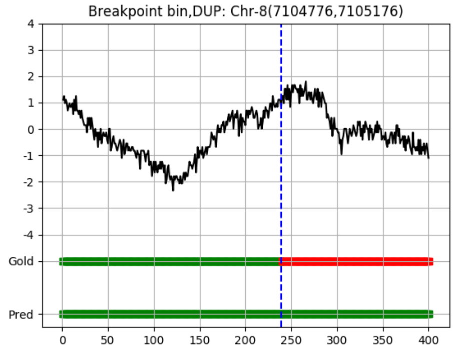

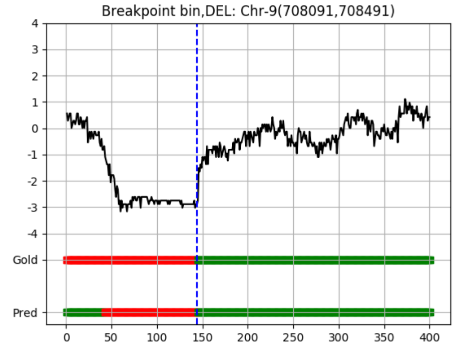

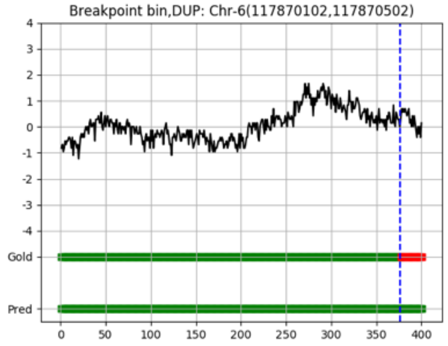

### Table S3: The mean and standard derivation of to-gold-distances of different ranges for real data in-sample evaluation

| | | | | | | | | | | | | | | | | | | | | |
|---|---|---|---|---|---|---|---|---|---|---|---|---|---|---|---|---|---|---|---|---|
| | | **mean to-gold-distance** | | | | | | | | | | **std to-gold-distance** | | | | | | | | |
| | | <5 | [5,10) | [10,20) | [20,50) | [50,100) | [100,200) | [200,500) | [500,1000) | >=1000 | | <5 | [5,10) | [10,20) | [20,50) | [50,100) | [100,200) | [200,500) | [500,1000) | >=1000 |
| **na12878** | / | 2.61 | 7.17 | 14.38 | 34.84 | 64.26 | 138.01 | 315.54 | 670.18 | 7364.78 | | 1.25 | 1.27 | 2.78 | 8.39 | 11.18 | 27.46 | 84.24 | 128.24 | 14794.04 |
| | CNN | 2.32 | 7.22 | 14.42 | 31.43 | 68.3 | 135.27 | 323.1 | 673.54 | 7250.29 | | 1.32 | 1.31 | 2.81 | 8.27 | 14.08 | 27.92 | 88.19 | 138.21 | 14643.51 |
| | Unet | 1.42 | 6.95 | 13.99 | 32.44 | 70.69 | 140.38 | 314.6 | 668.66 | 7383.95 | | 1.26 | 1.39 | 2.76 | 9.15 | 14.68 | 27.13 | 83.37 | 143.55 | 14782.42 |
| **na19238** | / | 2.5 | 7.14 | 14.94 | 35.43 | 65.46 | 141.07 | 331.91 | 684.02 | 4738.9 | | 1.36 | 1.32 | 2.81 | 8.44 | 11.58 | 30.53 | 100.65 | 137.68 | 8350.85 |
| | CNN | 2.42 | 7.08 | 14.08 | 32.54 | 69.63 | 139.36 | 318.67 | 703.58 | 4928.44 | | 1.27 | 1.41 | 2.87 | 8.54 | 13.14 | 28.21 | 91.73 | 146.83 | 8568.06 |
| | Unet | 1.39 | 6.73 | 14.14 | 32.07 | 71.84 | 140.89 | 302.6 | 690.13 | 4638.67 | | 1.18 | 1.41 | 2.73 | 8.69 | 14.37 | 28.95 | 89.61 | 146.9 | 8289.47 |
| **na19239** | / | 2 | 7.32 | 14.87 | 35.6 | 64.81 | 140.02 | 308.49 | 696 | 3592.76 | | 1.2 | 1.38 | 2.71 | 8.39 | 12.04 | 31.16 | 89.6 | 147.07 | 6264.21 |
| | CNN | 2.16 | 6.97 | 14.44 | 30.98 | 67.77 | 149.08 | 307.25 | 691.74 | 3498.3 | | 1.21 | 1.42 | 2.8 | 7.98 | 13.98 | 31.28 | 81.62 | 142.4 | 6172.19 |
| | Unet | 1.11 | 6.82 | 13.77 | 33.29 | 71.81 | 140.13 | 306.04 | 728.76 | 3690.94 | | 1.31 | 1.41 | 2.64 | 8.42 | 13.52 | 28.75 | 86.73 | 150.23 | 6363.64 |
| **HG002** | / | 2.12 | 7.2 | 14.55 | 34.1 | 62.53 | 143.1 | 302.61 | 759.22 | 2026.86 | | 1.38 | 1.37 | 2.88 | 8.51 | 11.58 | 28.34 | 77.88 | 128.36 | 1978.93 |
| | CNN | 2.1 | 6.88 | 14.14 | 30.98 | 67.3 | 142.33 | 295.96 | 767.19 | 2100.7 | | 1.33 | 1.44 | 2.87 | 8.35 | 13.42 | 28.08 | 77.41 | 126.57 | 2021.22 |
| | Unet | 0.85 | 6.73 | 13.49 | 33.19 | 68.85 | 143.94 | 287.65 | 753.48 | 2136.5 | | 1.29 | 1.48 | 2.62 | 9.14 | 13.06 | 29.82 | 74.57 | 126.1 | 2038.64 |

### Table S4: derivation of to-gold-distances of different ranges for real data cross-sample evaluation

| | | | | | | | | | | | | | | | | | | | | |
|---|---|---|---|---|---|---|---|---|---|---|---|---|---|---|---|---|---|---|---|---|
| | | **mean to-gold-distance** | | | | | | | | | | **std to-gold-distance** | | | | | | | | |
| | | <5 | [5,10) | [10,20) | [20,50) | [50,100) | [100,200) | [200,500) | [500,1000) | >=1000 | | <5 | [5,10) | [10,20) | [20,50) | [50,100) | [100,200) | [200,500) | [500,1000) | >=1000 |
| **NA19238** | / | 2.35 | 7.27 | 15 | 35.17 | 65.55 | 141.26 | 329.71 | 689.41 | 4553.32 | | 1.39 | 1.3 | 2.8 | 8.27 | 11.61 | 30.25 | 97.58 | 136.66 | 7581.48 |
| | CNN | 2.04 | 6.66 | 13.85 | 32.45 | 71.26 | 140.74 | 314.26 | 705.37 | 4709.07 | | 1.24 | 1.4 | 2.86 | 8.28 | 14.26 | 28.69 | 90.31 | 148.2 | 7933.48 |
| | Unet | 1.29 | 6.62 | 13.84 | 32.44 | 70.82 | 143.58 | 310.9 | 690.37 | 4493.61 | | 1.35 | 1.43 | 2.79 | 8.64 | 13.72 | 30.29 | 89.01 | 140.73 | 7707.92 |
| **NA19239** | / | 2.06 | 7.39 | 15.17 | 35.65 | 65.1 | 140.46 | 321.27 | 686.16 | 3923.15 | | 1.25 | 1.37 | 2.76 | 8.41 | 11.96 | 31.29 | 91.95 | 141.3 | 6207.5 |
| | CNN | 2.13 | 6.69 | 13.6 | 31.83 | 70.95 | 141.58 | 322.54 | 705.53 | 3974.33 | | 1.29 | 1.39 | 2.72 | 8.57 | 13.97 | 28.4 | 96.15 | 145.04 | 6249.28 |
| | Unet | 1.26 | 7.03 | 13.86 | 33.26 | 70.6 | 141.97 | 322.93 | 708.59 | 3932.57 | | 1.33 | 1.44 | 2.71 | 8.94 | 12.88 | 30.1 | 92.15 | 128.35 | 6207.13 |

**Table S5: Each repeat one result of 5-fold cross validation on the simulation data.**

| 80% train 20% test | | | | | | | | | |
|---|---|---|---|---|---|---|---|---|---|
| | RunID | AUC | Sensitivity | FDR | Precision | Recall | All-dice | BK-dice | all-IOU | BK-IOU |
| Unet | 1 | 0.8911 | 0.8619 | 0.0837 | 0.9163 | 0.8619 | 0.822 | 0.8503 | 0.6979 | 0.7397 |
| | 2 | 0.8945 | 0.8588 | 0.0741 | 0.9259 | 0.8588 | 0.8231 | 0.8489 | 0.6995 | 0.7376 |
| | 3 | 0.8951 | 0.8544 | 0.069 | 0.931 | 0.8544 | 0.8268 | 0.8487 | 0.7048 | 0.7372 |
| | 4 | 0.8957 | 0.8465 | 0.0604 | 0.9396 | 0.8465 | 0.8247 | 0.843 | 0.7018 | 0.7286 |
| | 5 | 0.8936 | 0.8513 | 0.0689 | 0.9311 | 0.8513 | 0.8232 | 0.8455 | 0.6996 | 0.7324 |
| | AVG | 0.894 | 0.85458 | 0.07122 | 0.92878 | 0.85458 | 0.82396 | 0.84728 | 0.70072 | 0.7351 |
| SVM | | 0.8661 | 0.8133 | 0.0898 | 0.9102 | 0.8133 | | | | |
| CNN | 1 | 0.8747 | 0.8428 | 0.0988 | 0.9012 | 0.8428 | 0.7957 | 0.8238 | 0.6607 | 0.7004 |
| | 2 | 0.8788 | 0.8531 | 0.0996 | 0.9004 | 0.8531 | 0.801 | 0.8286 | 0.6681 | 0.7074 |
| | 3 | 0.8751 | 0.859 | 0.111 | 0.889 | 0.859 | 0.7986 | 0.8293 | 0.6647 | 0.7083 |
| | 4 | 0.8751 | 0.8519 | 0.1055 | 0.8945 | 0.8519 | 0.7983 | 0.8281 | 0.6643 | 0.7066 |
| | 5 | 0.8758 | 0.8523 | 0.1042 | 0.8958 | 0.8523 | 0.7957 | 0.8252 | 0.6608 | 0.7025 |
| | AVG | 0.8759 | 0.85182 | 0.10382 | 0.89618 | 0.85182 | 0.79786 | 0.827 | 0.66372 | 0.70504 |

| 20% train 80% test | | | | | | | | | | |
|---|---|---|---|---|---|---|---|---|---|---|
| | | AUC | Sensitivity | FDR | Precision | Recall | All-dice | BK-dice | all-IOU | BK-IOU |
| Unet | 1 | 0.8817 | 0.8514 | 0.0922 | 0.9078 | 0.8514 | 0.7974 | 0.8247 | 0.6631 | 0.7019 |
| | 2 | 0.8844 | 0.8511 | 0.0866 | 0.9134 | 0.8511 | 0.8066 | 0.8343 | 0.676 | 0.7157 |
| | 3 | 0.8831 | 0.8476 | 0.0861 | 0.9139 | 0.8476 | 0.8033 | 0.8337 | 0.6712 | 0.7149 |
| | 4 | 0.8802 | 0.8336 | 0.078 | 0.922 | 0.8336 | 0.8037 | 0.8271 | 0.6719 | 0.7054 |
| | 5 | 0.883 | 0.8594 | 0.0967 | 0.9033 | 0.8594 | 0.8042 | 0.8356 | 0.6726 | 0.7178 |
| | AVG | 0.88248 | 0.84862 | 0.08792 | 0.91208 | 0.84862 | 0.80304 | 0.83108 | 0.67096 | 0.71114 |
| SVM | | 0.8576 | 0.8057 | 0.0998 | 0.9002 | 0.8057 | | | | |
| CNN | | 0.8432 | 0.8223 | 0.1394 | 0.8606 | 0.8223 | 0.7476 | 0.7853 | 0.5971 | 0.6467 |
| | | 0.8394 | 0.8363 | 0.1567 | 0.8433 | 0.8363 | 0.7469 | 0.7903 | 0.5964 | 0.6536 |
| | | 0.8438 | 0.8334 | 0.1461 | 0.8539 | 0.8334 | 0.7504 | 0.7925 | 0.6005 | 0.6565 |
| | | 0.8424 | 0.8172 | 0.1378 | 0.8622 | 0.8172 | 0.7454 | 0.7848 | 0.5942 | 0.646 |
| | | 0.8445 | 0.8237 | 0.1391 | 0.8609 | 0.8237 | 0.7489 | 0.7883 | 0.5987 | 0.6507 |
| | AVG | 0.84266 | 0.82658 | 0.14382 | 0.85618 | 0.82658 | 0.74784 | 0.78824 | 0.59738 | 0.6507 |
